# Quantifying charge state heterogeneity for proteins with multiple ionizable residues

**DOI:** 10.1101/2021.08.31.458420

**Authors:** Martin J. Fossat, Ammon E. Posey, Rohit V. Pappu

## Abstract

Ionizable residues can release and take up protons and this has an influence on protein structure and function. The extent of protonation is linked to the overall pH of the solution and the local environments of ionizable residues. Binding or unbinding of a single proton generates a distinct charge microstate defined by a specific pattern of charges. Accordingly, the overall partition function is a sum over all charge microstates and Boltzmann weights of all conformations associated with each of the charge microstates. This ensemble-of-ensembles description recast as a *q*-canonical ensemble allows us to analyze and interpret potentiometric titrations that provide information regarding net charge as a function of pH. In the *q*-canonical ensemble, charge microstates are grouped into mesostates where each mesostate is a collection of microstates of the same net charge. Here, we show that leveraging the structure of the *q*-canonical ensemble allows us to decouple contributions of net proton binding and release from proton arrangement and conformational considerations. Through application of the *q*-canonical formalism to analyze potentiometric measurements of net charge in proteins with repetitive patterns of Lys and Glu residues, we are able to determine the underlying mesostate pK_a_ values and, more importantly, we estimate relative mesostate populations as a function of pH. This is a strength of using the *q*-canonical approach and cannot be obtained using purely site-specific analyses. Overall, our work shows how measurements of charge equilibria, decoupled from measurements of conformational equilibria, and analyzed using the framework of the *q*-canonical ensemble, provide protein-specific quantitative descriptions of pH-dependent populations of mesostates. This method is of direct relevance for measuring and understanding how different charge states contribute to conformational, binding, and phase equilibria of proteins.

**STATEMENT OF SIGNIFICANCE:** The net charge of a protein in solution is governed by the overall pH as well as context and conformational contexts. Measurements of net charge are accessible via techniques such as potentiometry that quantify the buffering capacity of a protein solution. Here, we use the formal structure of the *q*-canonical ensemble to identify charge states that are compatible with a measured net charge profile as a function of pH. Our approach highlights how measurements of charge, decoupled from measurements of conformation, can be used to identify the ensembles of charge states that contribute to the overall population for given solution conditions. The methods introduced will be useful for measuring charge states and interpreting these measurements in different contexts.

## INTRODUCTION

Ionizable residues make key contributions to protein structure and function (1–6). They influence protein stability, solubility, and interactions mediated by the surfaces of the folded states of proteins (7–15), specifically in active sites of enzymes (16–18). Ionizable residues also feature prominently in intrinsically disordered proteins (IDPs) (19, 20). Several studies have documented the important contributions made by ionizable residues to conformational (21–26), binding (27-33), and phase equilibria of IDPs (34–42).

The charge states of ionizable residues can be regulated or altered through a variety of mechanisms that are collectively known as *charge regulation* (43–45). Active processes enable charge regulation through enzyme catalyzed reactions that enable post-translational modifications (46) such as lysine acetylation (47), arginine citrullination (48), serine / threonine / tyrosine phosphorylation (49–51), and deamidation of asparagine / glutamine (52). In addition to enzyme catalyzed charge regulation, the charge states of ionizable residues can also be altered by spontaneous processes such as binding or release of protons (2, 3, 7, 53-59), and the preferential accumulation or exclusion of solution ions around regions of high charge density (60–67).

Charge regulation via proton uptake or release is influenced by a combination of sequence contexts, solution conditions, and the linked effects of conformational equilibria (68, 69). Importantly, multiple charge states are accessible depending on solution conditions. As a result, each protein sequence is an ensemble of charge microstates, with each microstate being a distinct sequence defined by the charge states of ionizable residues. We recently introduced the *q*-canonical ensemble to describe ensembles of charge microstates *and* conformations associated with each of the microstates (70). The structure of the *q*-canonical ensemble, see **Figure 1**, is as follows: We consider a protein sequence comprising *n_q_* ionizable residues, *N*_fixed_ atoms that do not change with protonation / deprotonation, and a net charge of *q* when 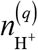 protons are bound to the protein sequence. If there are 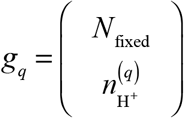 ways in which 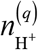 protons can be bound, then there are *g_q_* distinct charge microstates for a net charge of *q*. At temperature *T*, in a solution volume *V*, each charge microstate *i* with net charge *q* will access a canonical ensemble of conformations. For this ensemble, the partition function is written as:

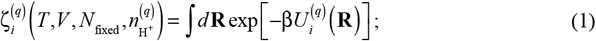

**Figure 1:**
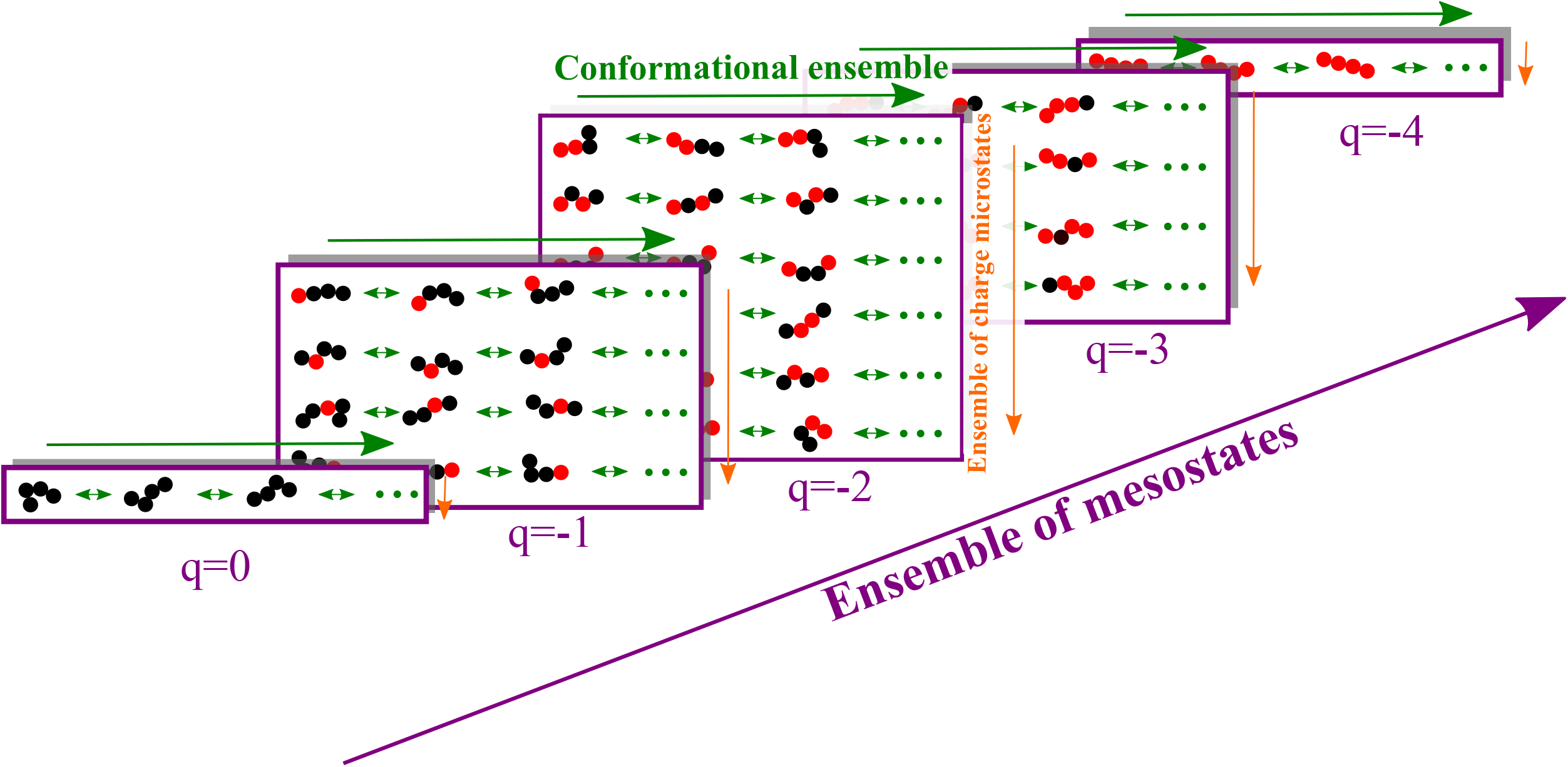
Structure of the *q*-canonical ensemble. Schematic depicting the structure of the *q*-canonical ensemble. The ensemble of mesostates (purple arrow) encompasses all mesostates (purple rectangles). Each mesostate encompasses an ensemble of charge microstates (orange arrows) corresponding to the different ways residue in their charged state (red circles) can be arranged with respect to those in their uncharged state (black circles). Finally, each microstate has a distinct conformational ensemble (green arrows).

Summing over the *g_q_* ways in which when 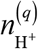 protons can be bound leads to a sum over canonical partition functions written as:

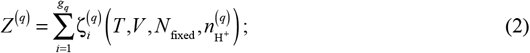

The partition function, written as the sum over the different numbers 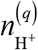 of protons can be bound becomes:

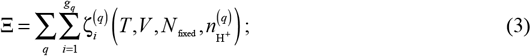

The states summed over in equation (3) are the set of conformations accessible for each charge microstate, with the outer two summations running over all possible microstates. The partition function Ξ describes the *q*-canonical ensemble. Next, we rewrite Ξ to account for the chemical potential of the proton, which is defined by the pH of the solution. Accounting for pH, Ξ becomes:

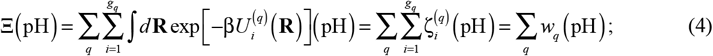

The conformational ensemble for each charge microstate contributes to 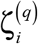, and the sum over all microstates with a charge *q*, which we refer to as a charge mesostate, contributes to *W_q._* A mesostate is made up of all charge microstates that have the same net charge, and the sum over *q* is a sum over all mesostates. Accordingly, we rewrite the sum over *q* as a sum over mesostates such that the probability that the protein of interest will have a charge *q* at a given pH can be written as *p_q_*, and this is computed as:

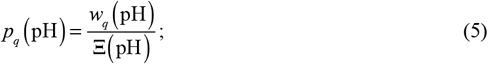

Given this formalism, the net charge as a function of pH can be written as:

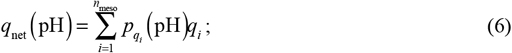

Note that *p_q_i__* (pH) is the probability associated with mesostate *i* that has charge *q_i_*. Potentiometric measurements provide a direct readout of net charge as a function of pH. Here, we present a new approach to analyzing potentiometric data that is based on using equation (6) to describe these data. Our approach relies on the structure of the *q*-canonical ensemble, which allows for a formal decoupling of measurements of charge from measurements of structure. This is achieved by estimating the mesostate weights *p_q_i__* (pH) from potentiometric measurements using a Monte Carlo based fitting procedure. This information can be subsequently combined with simulations as well as measurements of conformational averages or populations, performed as a function of pH, to identify the microstates that contribute most significantly to a mesostate. The key point is that subsequent computations of microstate probabilities and conformational distributions can be constrained by the values for *p_q_i__* (pH) that we obtain using potentiometric measurements.

In the following sections, we summarize the challenges posed by acid-base equilibria of proteins with multiple ionizable residues. Such systems are referred to as polyacids (54, 71-74). We then describe how the *q*-canonical formalism can be interfaced with potentiometric measurements (75–77). This is followed by a presentation of our results from application of the *q*-canonical formalism to analyze potentiometry data for three model systems that are repeats of Lys and Glu. We conclude with a discussion of the broader implications of our findings, the general use of our approach, and the insights that should be forthcoming through joint use of the *q*-canonical ensemble, conformational sampling, and separate measurements of proton binding and conformational equilibria.

## THEORY

### Acid-base equilibria for polyacids

For a sequence with *n* ionizable residues, there are, in theory, 2^*n*^ distinct charge microstates to consider. However, for sequences that are mixtures of acids and bases, the number of thermodynamically relevant charge microstates can be significantly reduced by eliminating forbidden states (70). Any microstate that involves the coexistence of neutral versions of acidic and basic residues can, to first approximation, be ignored as forbidden microstates. This is because the intrinsic pK_a_ values are such that the protonation of acids and deprotonation of bases are unlikely to occur simultaneously. Classification of microstates as being forbidden and ignoring these in subsequent calculations does not materially alter the estimated partition function (70). Following the pruning of forbidden microstates, the number of thermodynamically relevant charge microstates is designated as Ω_micro_. Even with pruning to eliminate forbidden microstates, Ω_micro_ will increase exponentially with the number of ionizable residues *n* (**Figure S1**). Comparative descriptions of acid-base equilibria for simple systems vs. polyacids shows how the large number of charge microstates must be accounted for when assessing pH-dependent titrations for polyacids.

For a monoacid such as a single glutamic acid, the pK_a_ is defined for the reaction e ⇌ E + H^+^. Here, e and E, respectively denote the protonated and deprotonated versions of Glu, and H^+^ refers to the proton that is released upon deprotonation. The standard state ionization free energy is: 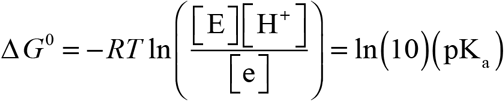. The picture becomes more complicated for polyacids because of the increase in the number of reactions that involve the loss of a proton (78). To illustrate this point, we consider the case of ac-(Glu)_2_-nme, where ac and nme, respectively refer to N-acetyl and N’-methylamide. There are four separate reactions that involve the loss of a single proton. These are:

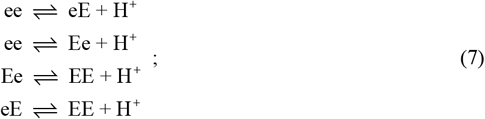

Each of the reactions shown in Equation (7) can be assigned a pK_a_. We refer to these as primary pK_a_ values, as they involve the loss of exactly one mole of protons from a single site for each mole of advancement of the reaction. For a system with *n* ionizable residues, there will be a set of *n*2^*n-1*^ primary pK_a_ values. These cannot be measured, although they can be approximated (79–81).

A mesostate is made up of all charge microstates that have the same net charge – see Equations (4) and (5). Each mesostate is assigned a label *q_k_* where *q* denotes the net charge associated with each microstate within the mesostate and *K* denotes the total number of microstates within the mesostate. For example, the designation of + 1_3_ implies that there are three microstates in the mesostate, and all these microstates have a net charge of + 1. The total number of mesostates, denoted as *N*_meso_, scales linearly with the number of ionizable residues (**Figure S2**). We can rewrite the acid-base equilibria in Equation (7) using the notation for mesostates (**Figure 2**) and rewrite the relevant reactions for the release of one mole of protons as follows:

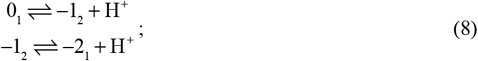

**Figure 2:**
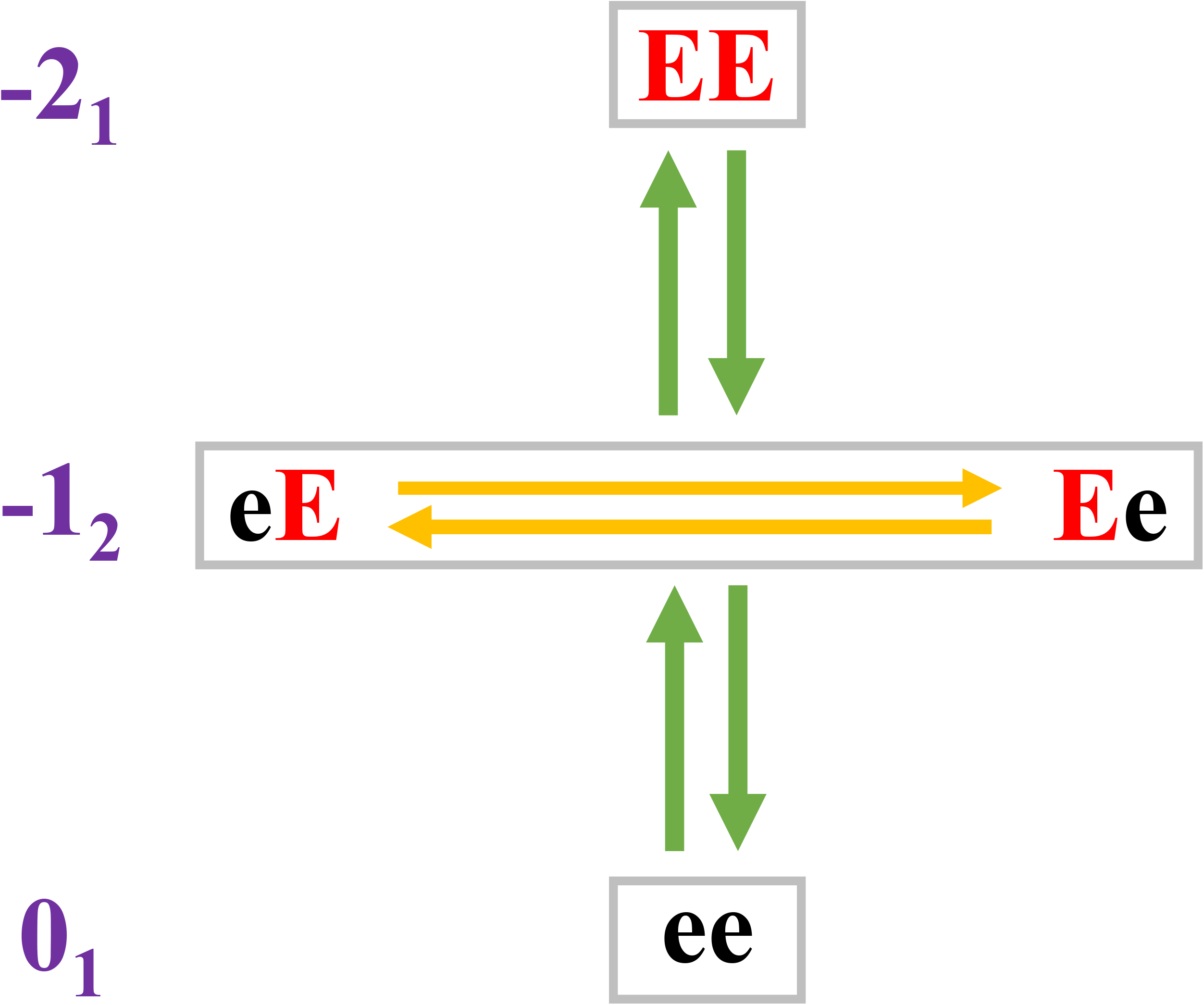
Schematic showing charge microstates and the grouping into mesostates for ac-(Glu)_2_-nme. Here ace and nme refer to N-acetyl and N’-methylamide, respectively. Protonated and deprotonated Glu residues are depicted as e and E, respectively. The charge microstates ee, eE, Ee, and EE can be grouped into mesostates, and the designation of mesostates, shown in *q_k_* format is shown in purple.

Here, (0_1_,–1_2_) and (–1_2_,–2_1_) are two pairs of adjacent mesostates. Adjacency refers to mesostates whose net charge differs by ±1*e*. Each reaction in Equation (8) can also be assigned a pK_a_ value. These mesostate pK_a_ values are different from the set of primary pK_a_ values. For *N*_meso_ mesostates, there will be *N*_meso_ – 1 mesostate pK_a_ values corresponding to transitions between pairs of adjacent mesostates. While the set of primary pK_a_ values cannot be measured, the set of mesostate pK_a_ values can be inferred from measurements of net charge as a function of pH, providing we are able to extract the pH-dependent populations of mesostates – a problem we solve here by leveraging the structure of the *q*-canonical ensemble.

For the case of the ac-(Glu)_2_-nme system, there are at least two distinct scenarios that could correspond to a seemingly reasonable definition of unshifted pK_a_ values. According to one assumption, the pK_a_ for a transition between two mesostates would be identical to the intrinsic pK_a_. This would imply that Δ*G*_*q_k_*→(*q*-1)_*l*__ = Δ*G*_e → E_. However, this assumption ignores the number of reactions within a single mesostate, each involving a distinct microstate, that can contribute to the loss of a proton. The second model for unshifted pK_a_ values corresponds to the assumption that: Δ*G*_ee → eE_ = Δ*G*_e → E_. This model assumes that the primary pK_a_ values can only be assigned to reactions involving individual microstates rather than collections of microstates. Here, the unshifted pK_a_ associated with transitions between mesostates can be derived through proper accounting of the diversity of charge microstates per mesostate. We illustrate these points using reactions for the ac-(Glu)_2_-nme system. The mesostate –1_2_ includes the microstates Ee and eE. Therefore, the concentration of mesostate –1_2_, denoted as [–1_2_] is written as: [–1_2_] = [Ee] + [eE]. Accordingly, the equilibrium constant for dissociation of a mole of protons from mesostate –1_2_ is written as:

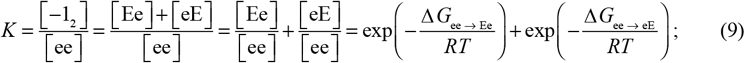

The free energy of ionization of a single mesostate, for the specific example considered here, may be written as:

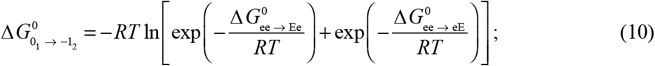

If we assume that primary pK_a_ values correspond to those of model compounds, we can use Equation (10) for polyacids by accounting for the number of microstates per each mesostate. The free energy of ionization that transforms mesostate *q_k_* to (*q*–1)_*l*_ of a polyacid, through the release of one mole of protons, may be written as:

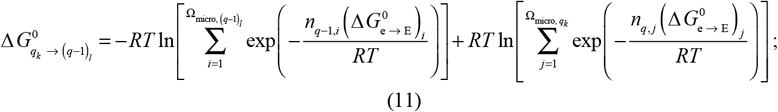

Here, Ω_micro,*qk*_ and Ω_micro, (*q*-1)_*l*__ refer to the number of charge microstates within mesostates *q_k_* and (*q*–1)_*l*_, respectively; 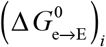 and 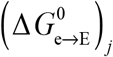. are the context dependent free energies of ionization; *n*_*q*-1,*i*_ and *n_q,j_* are the numbers of ionizable residues within microstates *i* and *j* that are in their basic forms. For example, *n_q_* = 1 for microstates eE, eEe, kK, KKk, etc. Likewise, *n_q_* = 2 for microstates eEE, kkK, etc. Here, k is the deprotonated version of lysine (K). For the transformation of mesostate 0_1_ to mesostate –1_2_, *n*_*q*-1,*i*_ = 1 for each of the microstates Ee and eE, whereas *n_q,j_* = 0 for the microstate ee.

Equation (11) reduces to Equation (10) if we set 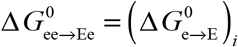 and 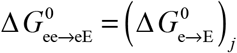. If all microstates within a mesostate are equiprobable, implying that the ionization free energy is independent of sequence context, then Equation (11) reduces to:

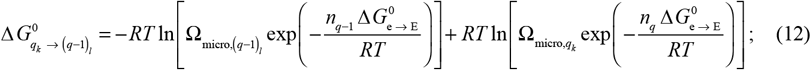

Upon further rearrangement, Equation (12) becomes:

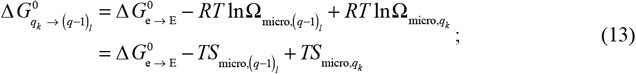

Equation (13) highlights the importance of microstate entropy that features in two terms: *S*_micro, *q_k_*_ = *R*lnΩ_micro, *q_k_*_ and *S*_micro, (*q*-1)_*l*__ = *R*lnΩ_micro,(*q*-1)_*l*__. These are the microstate entropies of mesostates *q_k_* and (*q*—1)_*l*_, respectively. The implication is that even for the simple scenario of context-independent pK_a_ values, the free energy difference between a pair of adjacent mesostates, which is a measure of the pK_a_ associated with the transition between a pair of adjacent mesostates, must account for the contributions of different microstate entropies for each mesostate. In general, the microstate weights will be non-identical. In this scenario, the pH-independent standard state free energy change associated with transformation from mesostate *q_k_* to (*q*—1)_*l*_ is written as:

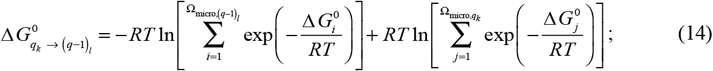

Here, 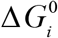 is the standard state free energy of microstate *i*. In the following sections we show how potentiometry, which measures the buffering capacity of a solute as a function of pH, can be used in conjunction with Equation (14), to derive mesostate populations as a function of pH.

### Extracting mesostate populations from potentiometry measurements

We use potentiometry to estimate the net charge of a protein as a function of pH. This is achieved by titrating the solution pH using a strong acid or a strong base. The buffering capacity of a protein solution, referenced to the buffering capacity of the protein-free solution, is used to estimate the profile of net charge vs. pH. Given the ensemble-averaged net charge vs. pH as an input, we can fit the measured profile to extract the pH-dependent population of mesostates. This information allows us to extract the contributions made by distinct mesostates to the pH-dependent net charge profile. The measured net charge *q*_net_ as a function of pH is given in Equation (6), where *q_i_* is the charge associated with mesostate *i* and *p_i_*(pH) is the probability of populating mesostate *i* for a given pH. Given knowledge of *q*_net_(pH), and the charge associated with each mesostate, we can estimate the pH-dependent mesostate probabilities by fitting the measured profiles to Equation (9). The fitting procedure is based on a Metropolis Monte Carlo method (82) that minimizes the deviation between the measured and estimated charge profile by minimizing a cost function defined as:

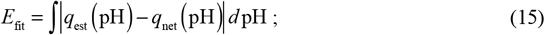

We choose absolute values rather than signed values to keep the cost function bounded between 0 and a positive number. In Equation (15), *q*_est_ and *q*_net_ are the estimated and measured net charge values for a given pH. The fitting procedure, summarized in **Figure 3**, can be seeded by assigning mesostate probabilities using intrinsic pK_a_ values, although as discussed in the Supporting Material, this is not essential. We use 10.7 and 4.34 for intrinsic pK_a_ values of Lys and Glu, respectively (83). The parameter Δ_max_ (see **Figure 3**) is set so that the maximal change in relative free energies of states i and j that are being perturbed will be no more than 0.01 kcal / mol. The Metropolis criterion for accepting or rejecting a proposed change in mesostate probabilities is of the form: 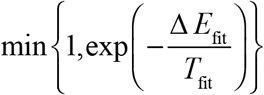. Here, Δ*E*_fit_ quantifies how the cost function *E*_fit_ has changed with the newly proposed mesostate probabilities. The parameter *T*_fit_ is a scalar parameter, which is a fictitious and unitless temperature that is chosen to enable efficient convergence of the Monte Carlo based fitting procedure. The optimal temperature will be system-specific, and is chosen by inspecting the acceptance ratio and making sure it is not close to one or below a numerical tolerance. Data from potentiometry measurements (see below) converted into profiles of net charge vs. pH are used to quantify how the mesostate probabilities evolve as a function of pH.

**Figure 3:**
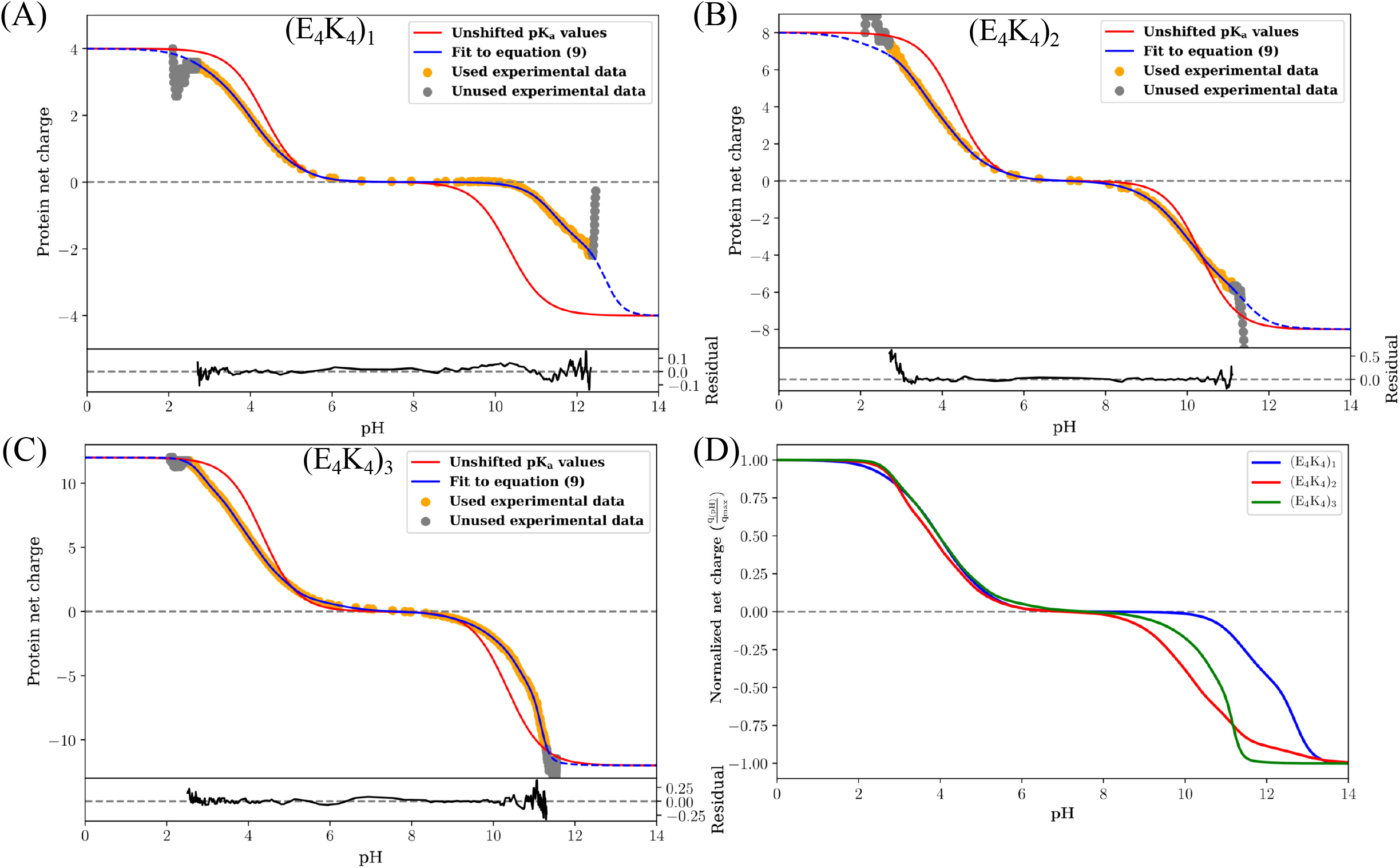
Flowchart summarizing the workflow for fitting data from potentiometric titrations to Equation (9).

Smoothing of the raw data was performed using a Savitzky-Golay algorithm (84) as implemented in the SciPy python package (https://scipy.github.io). The prescribed output resolution was set to 100 points per pH unit, with a window size of 101, and a second-order polynomial. Results of the smoothing procedure are shown in **Figure S3**. To enhance sampling efficiency, especially in the first steps of the fit, we introduce a drift correction, which returns the fitting parameters to the parameters of the best fit found to that point, if the current fit has a cost function of more than 125% of the best fit cost function. The best fit as a function of step number is depicted for two different priors in Movie S1 (all pK_a_ values set to 7) and Movie S2 (pK_a_ values set to their intrinsic pK_a_ values). Each Monte Carlo fitting procedure comprises 150000 steps. For each construct, we performed a total of 10 independent runs. The best fits obtained in each run are depicted as the blue curves of **Figure S5**. These fits show clear convergence, especially for the neutral pH region

It is worth emphasizing that the fitting procedure generates estimates for mesostate populations as a function of pH. Further parsing of these mesostate specific weights to unearth the contributions of mesostate-specific microstates will require additional information in the form of the full *q*-canonical Monte Carlo simulation that assesses the contributions of microstates and their conformations. These simulations can be constrained by the mesostate populations we derive using the procedures summarized in **Figure 3**.

## MATERIALS AND METHODS

### Reagents and peptides

Potassium chloride (KCl), hydrochloric acid (HCl), potassium hydroxide (KOH), and potassium hydrogen phthalate (KHP) were purchased from Sigma. We assessed the purity of KOH using pH measurements, which confirm that contaminants, if present are miniscule and do not affect the potentiometric titrations. Peptides were purchased from GenScript at >95% purity as HCl salts with acetylated N-termini and amidated C-termini. Salt content was determined by mass spectrometry. The identities of the peptides were confirmed using mass spectrograms provided by the vendor. Peptides were stored in lyophilized form at –20 °C in sealed containers in the presence of desiccant until they were used for experiments. We performed measurements for three peptides, excised from proteins with single alpha helical domains. The amino acid sequences of the three peptides were as follows: Ac-G-(E_4_K_4_)_n_-GW-NH_2_, where n = 1, 2, or 3. The peptides were capped at the N- and C-termini using N-acetyl (ace) and an amide, respectively. This avoids confounding effects from charged termini. Trp was used to enable precise measurements of concentration.

### Sample preparation

A sealed flask of ultrapure water was depleted of CO_2_ using a custom-built Schlenk line to apply alternating cycles of nitrogen gas and vacuum. The resulting CO_2_ depleted water and all solutions and samples prepared from this water were kept in sealed flasks or vials and stored in a glove bag (AtmosBag, Millipore-Sigma Z530204) filled with nitrogen gas. To further decrease the concentration of dissolved CO_2_, solutions prepared with the CO_2_-purged water were allowed to equilibrate with the nitrogen-filled (CO_2_-depleted) atmosphere in the glove bag prior to sealing the flask or vial. All transfers of these liquids between flasks or vials were carried out within the nitrogen-filled glove bag or using gas-tight Hamilton syringes to access sealed vials without exposing the contents to air. A stock solution of background solvent was prepared by dissolving KCl in CO_2_-depleted ultrapure water to which HCl had been added. The final solution contained 10 mM KCl and 5 mM HCl. This solution was further depleted of CO_2_ using the Schlenk line, after which it was sealed and stored in a nitrogen-filled glove bag until further use.

A 50 mM KOH solution was used as the titrant for all potentiometry experiments. A 200 mM KOH stock solution was prepared in the glove bag using CO_2_-depleted ultrapure water. This stock was further diluted to 50 mM KOH with CO_2_-depleted ultrapure water and sealed in a glass vial with a Teflon-coated rubber septum while in the nitrogen-filled glove bag. Peptides were dissolved in the CO_2_-depleted background solution at a concentration of ~200 μM. The final peptide concentrations used were 215 μM for E_4_K_4_, 95 μM for (E_4_K_4_)_2_, and 180 μM for (E_4_K_4_)_3_. Immediately prior to each potentiometry experiment, the peptide concentration was measured by UV/visible spectroscopy using Trp absorbance at 280 nm and an extinction coefficient of 5500 M^-1^ cm^-1^. A KHP solution at a concentration of 1 mM was prepared by massing a quantity of KHP powder into a glass vial using a precision mass balance, transferring the vial to the nitrogen-filled glove bag, and then adding CO_2_-depleted ultrapure water directly to the vial to dissolve the material. The sample was stored in a sealed vial in the nitrogen-filled glove bag until further use.

### Potentiometry

All measurements were carried out in a custom sample vial consisting of a glass screw-cap vial sealed with Teflon-coated rubber septum with a 5 mm hole (created with a biopsy punch) that accommodated the pH probe while still forming a seal. Titrant was delivered to the sealed vial through the rubber septum using a precision gas-tight μL syringe fitted with a repeating dispenser (Hamilton). Potentiometric measurements were carried out on an Orion Star A215 potentiometer (Thermo Scientific) using a pHT-micro combination probe with a platinum / Silamid double junction (YSI). A 2 mL sample was titrated with 50 mM KOH in 2 μL increments in all cases.

The concentration of the KOH solution was verified by titrating a known concentration of potassium hydrogen phthalate (KHP) having a well-documented pKa of 5.4, as follows: a 2 mL volume of KHP solution was titrated with ~50 mM KOH solution in 2 μL increments, and the pH of the solution was measured after each addition. A titration of 2 mL of background solution was carried out in the same manner. The resulting potentiometry curves (KHP and background) were analyzed by Gran analysis (85, 86) as well as by first and second derivative analyses to determine the precise KOH concentration (87, 88).

For each peptide, a 2 mL volume of peptide solution was titrated with ~50 mM KOH solution in 2 μL increments, and the pH of the solution was measured after each addition. A titration of 2 mL of background solution was carried out in the same manner. For each peptide titration, the KOH solution used as titrant was prepared fresh and calibrated with a KHP titration as described above.

A stable pH reading is essential for accurate potentiometric measurements, and this can be difficult to achieve under conditions of low solute concentration. For example, it is extremely difficult, if not impossible to obtain a stable pH reading in a sample of ultrapure water due to the lack of buffering capacity and the lack of salt ions in the sample. In our experiments, we found that 10 mM salt was sufficient to ensure signal stability, as evidenced by successful measurement of stable pH in background titrations of 10 mM KCl and 5 mM HCl with no peptide (**Figures S8-S10**). In peptide-containing samples, the presence of buffering moieties at millimolar concentrations (1.72 mM for E_4_K_4_, 1.52 mM for (E_4_K_4_)_2_, and 4.32 mM for (E_4_K_4_)_3_) provided further stability to pH readings. Additional measurement stability was facilitated by the use of a pH probe with a platinum wire junction, which allows optimum electrolyte flow, and by allowing the signal to plateau and stabilize for a minimum of 50 seconds (longer near neutral pH) for each data point.

### Calculation of peptide net charge

Since KOH is a strong base, we can assume that for each mole of OH added, one mole of protons is consumed, resulting in water as the product. We use the known concentration of the KOH solution to calculate the number of moles of KOH from the volume of KOH added. The difference in KOH volume added between the titration curves for peptide and background represents a quantity that is proportional to the number of moles of protons introduced by the peptide. Since the peptide concentration is known, we can calculate the number of protons per peptide. The number of protons released over the course of the titration is equivalent to the change in net charge of the peptide. Therefore, the net charge of the peptide can be determined as a function of pH if the net charge is known at the starting point of the titration. Since the titration is started at a low pH (~2.0) that is below the pK_a_ values of all ionizable groups present, we can assume that all ionizable groups are protonated, and the net charge on the peptide should be equal to the number of cationic residues, (Lys in the case of (E_4_K_4_)_n_ peptides).

Following the approach of Nozaki and Tanford (89), the data were transposed to facilitate comparison of the volume of KOH added in the background titration vs. the peptide titration at each pH value (2, 90). A linear interpolation between each data point within a given titration curve was carried out so that the background curve could be subtracted from the peptide curve at any pH value. The resulting quantities represent the difference in KOH volume added in the background titration vs. the peptide titration at a given pH value. Multiplying these values by the known concentration of KOH gives the difference in number of moles of KOH between the two curves, and this is equivalent to the number of protons introduced by the peptide. Dividing this value by the number of moles of peptide (calculated as peptide concentration multiplied by sample volume) quantifies the moles of protons per moles of peptide, which is equivalent to relative change in net charge (**Figures S11-S13**).

As explained above, for (E_4_K_4_)_n_ peptides, the number of Lys residues provides the offset value needed to shift the calculated moles of protons per peptide to the appropriate register for reporting the net charge of the peptide. In practice, there are a few sources of uncertainty that may require adjustments to both the scaling and the offset of the net charge curve. These include any uncertainties in measurements of concentration, and the presence of trace amounts of residual acid in the peptide sample. To account for these errors, we scale the protein net charge curve such that the inflection point around neutral pH has a charge of 0. This is the most parsimonious rescaling, since any other rescaling would lead to mesostates with non-zero charge being dominant population around pH 7.0, implying that the 0_1_ mesostate would never be dominant in the ensemble. The rescaling factor used for each construct is 0.92, 0.76 and 0.79 for (E_4_K_4_)_1_, (E_4_K_4_)_2_, and (E_4_K_4_)_3_, respectively.

## RESULTS

Sequences with a consensus repeat consisting of (E_4_X_4_)_n_ or (E_3_X_3_)_n_ where X is either an Arg or Lys have been shown to form alpha helical conformations (91–94). These single alpha helix forming sequences belong to larger proteins (93). Helicity is known to increase with the repeat length (91). The single alpha helices are thought to be useful in force transduction in myosin (92) and have been deployed as spectroscopic rulers for Förster resonance energy transfer measurements, *in vitro* and in cells (95). The overall helicity has been ascribed to a network of intra-helical salt bridges between residues i and i+4 (91, 94).

Here, we ask if a single mesostate *viz*., 0_1_ dominates over the entire accessible pH range, irrespective of the number of repeats. This question is motivated by observations showing a remarkable stability of the alpha helical conformations across a wide range of pH values (94, 96). We reasoned that it is unlikely for networks of salt bridges to be operative under conditions where Glu is protonated, or Lys is deprotonated. A definitive assessment of the curious pH insensitivity requires knowledge of how the net charge varies with pH, and we measure this directly. We can then use these data, in conjunction with the *q*-canonical ensemble to infer the populations of each of the relevant mesostates as a function of pH. The measurements reveal a series of insights that are summarized next.

### Fitting of potentiometric data

**Figures 4A, 4B, and 4C** show the quality of the fits obtained for each of the three peptides. Further, Movie S1 provides a direct visualization of how the fitting procedure evolves. In each panel of **Figure 4**, the gray circles denote the raw experimental data that are not used in the fitting procedure. The orange points are the raw data used for the fitting procedure. The fitting procedure was applied to the smoothed version of the data to have a uniform resolution across the pH range to ensure that evaluation of the cost function is uniformly distributed. The raw data and comparisons to the smoothed data are shown in **Figure S3**. Fits to the data are shown using two models, depicted using blue vs. red curves. The red curves are charge profiles calculated by assuming model compound pK_a_ values for all the ionizable residues, although the microstate entropies per mesostate are different. In this model, the only parameters that distinguish different mesostates are net charge and the microstate entropy per mesostate. These fits, which do not describe the data well, clearly indicate that there are shifted pK_a_ values in each of the three sequences. The blue curves show the quality of fit obtained through the Monte Carlo fitting procedure that uses Equation (16). The quality of the fits is shown in terms of residuals included at the bottom of each panel. The mesostate probabilities change with respect to the unshifted model, and these changes are reflected in the improved fits to the net charge vs. pH profiles. We further assessed the quality of the fits by calculating the pH-dependent derivatives of the net charge profiles, for the experimental data and for the numerical fits. These comparisons are shown in **Figure S4. Figure 4D** shows a comparison of the normalized net charge vs. pH for the three peptides. If the unshifted model were to be valid, then the three curves should collapse on one another. Clearly, this is not the case, and it points to a sequence- and context-dependent contribution from the increasing diversity of mesostates with peptide length.

**Figure 4:**
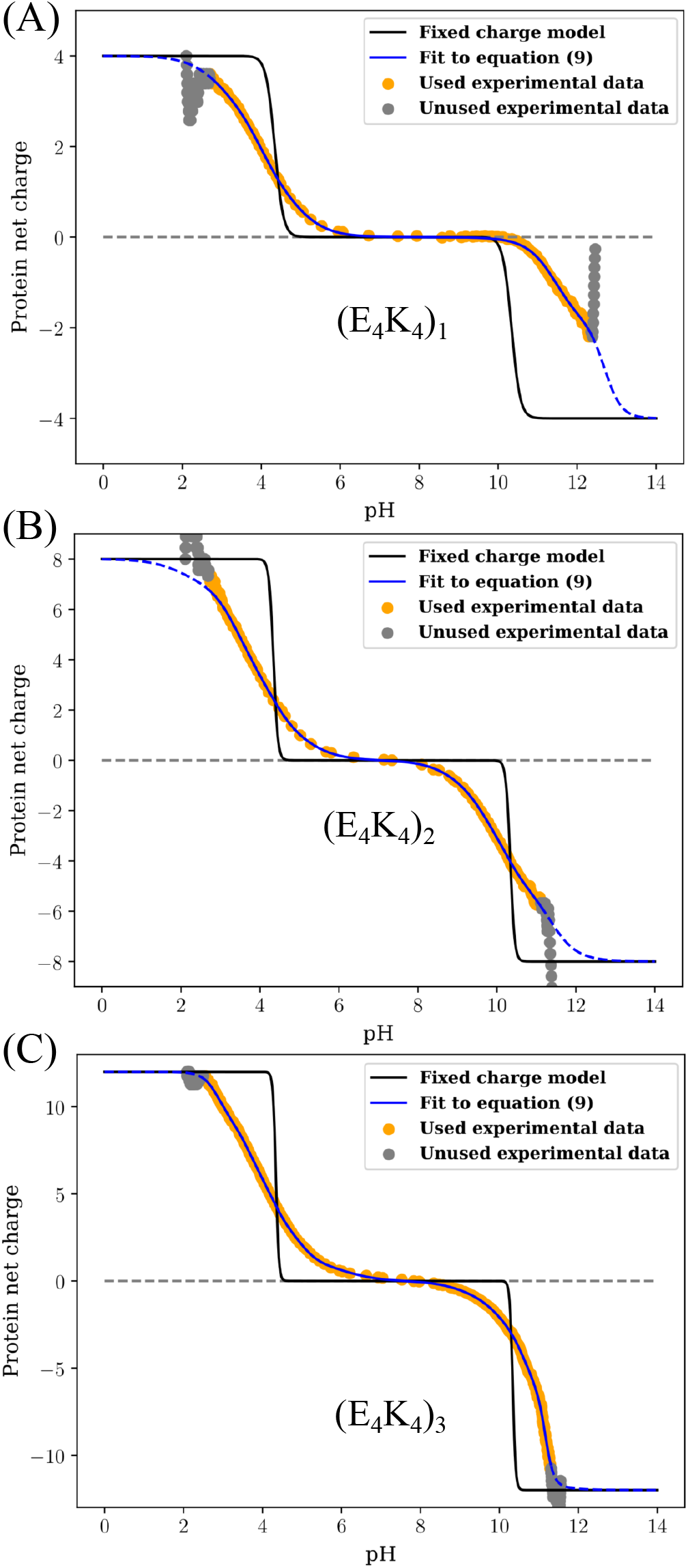
Raw data and results obtained from fitting the model based on unshifted pK_a_ values vs. the full model – Equation (9) – to the profile of net charge vs. pH obtained from potentiometric titrations. Panel (D) shows a comparison of the fitted profiles for the three peptides. To enable these comparisons, the ordinate is rescaled to be normalized by the magnitude of the maximal charge realizable for each system.

We also compared charge profiles calculated using a fixed charge model to the measured profiles from potentiometry. Here, we assume that Lys is always protonated below its intrinsic pK_a_ and Glu is always deprotonated above its intrinsic pK_a_. The results are shown in the three panels of **Figure 5**. Again, each panel includes the gray circles, orange points, and blue curves that are identical to those in **Figure 4**. Clearly, the fixed charge model does not recapitulate how the measured charge varies with pH, at least for the (E_4_K_4_)_n_ series of peptides.

**Figure 5:**
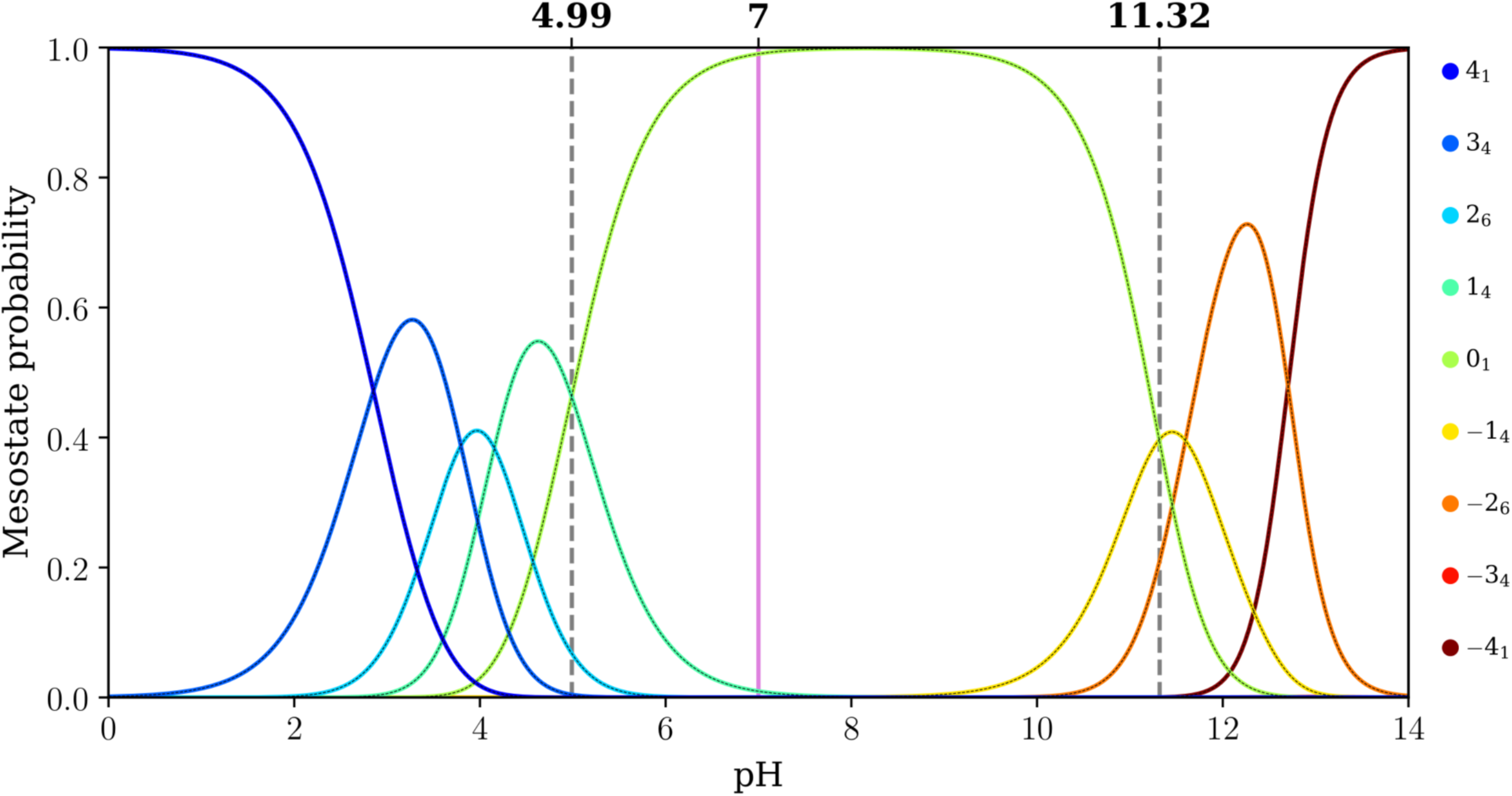
Comparison of the raw data, fits obtained using the full model, and a model that uses fixed charges. The results are shown for each of the three peptides. In each panel, the black curve corresponds to the fixed charge model. This model does a poor job of recapitulating the titration curve.

### Interpretations from the measured charge profiles

The profile for (E_4_K_4_) shows a plateau in the pH range between 7 and 9 (**Figure 4a**). This observation is confirmed by calculations of the derivative as a function of pH, which show that the derivative is essentially zero in the pH range between 7 and 9 (**Figure S4a**). The implication is that none of the acids or bases are titrating their charge states in this pH range. Therefore, a single mesostate must be dominant in this pH range for the E_4_K_4_ peptide. It is noteworthy, however, that the plateau region of the blue curve in **Figure 4a** is larger than that of the red curve. The latter corresponds to the model that uses unshifted, model compound pK_a_ values while accounting for differences in microstate entropies of the different mesostates. Thus, the data indicate a downshift vs. an upshift in the pK_a_ values for the acids and the bases, respectively. In contrast to the (E_4_K_4_) peptide, the two longer peptides (E_4_K_4_)_2_ and (E_4_K_4_)_3_ do not have a pH range where the charge profile plateaus (**Figures 4b** and **4c**), and this is confirmed by the absence of a pH region where the derivative is zero (**Figures S4b** and **S4c**). This suggests that at every pH, there is at least more than one mesostate (and hence microstate) that contributes to the measured charge profiles, especially for (E_4_K_4_)_2_.

### Insights from pH-dependent mesostate populations

Fitting of the data for net charge vs. pH using the *q*-canonical ensemble formalism yields the pH-dependence of mesostate populations. The results are shown in **Figures 6-8** for E_4_K_4_, (E_4_K_4_)_2_, and (E_4_K_4_)_3_, respectively. The dashed vertical lines in **Figure 6** help quantify mesostate pK_a_ values. For E_4_K_4_, the +1_4_ and 0_1_ mesostates have equal likelihoods of being populated at a pH of 4.99. This implies that the pK_a_ associated with the transition between +1_4_ and 0_1_ mesostates is 4.99. Using a similar approach, we find that the pK_a_ value associated with the transition between 0_1_ and −1_8_ mesostates is 11.32. Therefore, the 0_1_ mesostate is dominant within the pH range between 4.99 and 11.32, with the gap between the two mesostate pK_a_ values being 6.33 pH units. However, the dominance of the 0_1_ mesostate weakens with increasing numbers of E_4_K_4_ repeats, and the prominence of the mesostates with net charges of ±1 increases with the number of E_4_K_4_ repeats.

**Figure 6:**
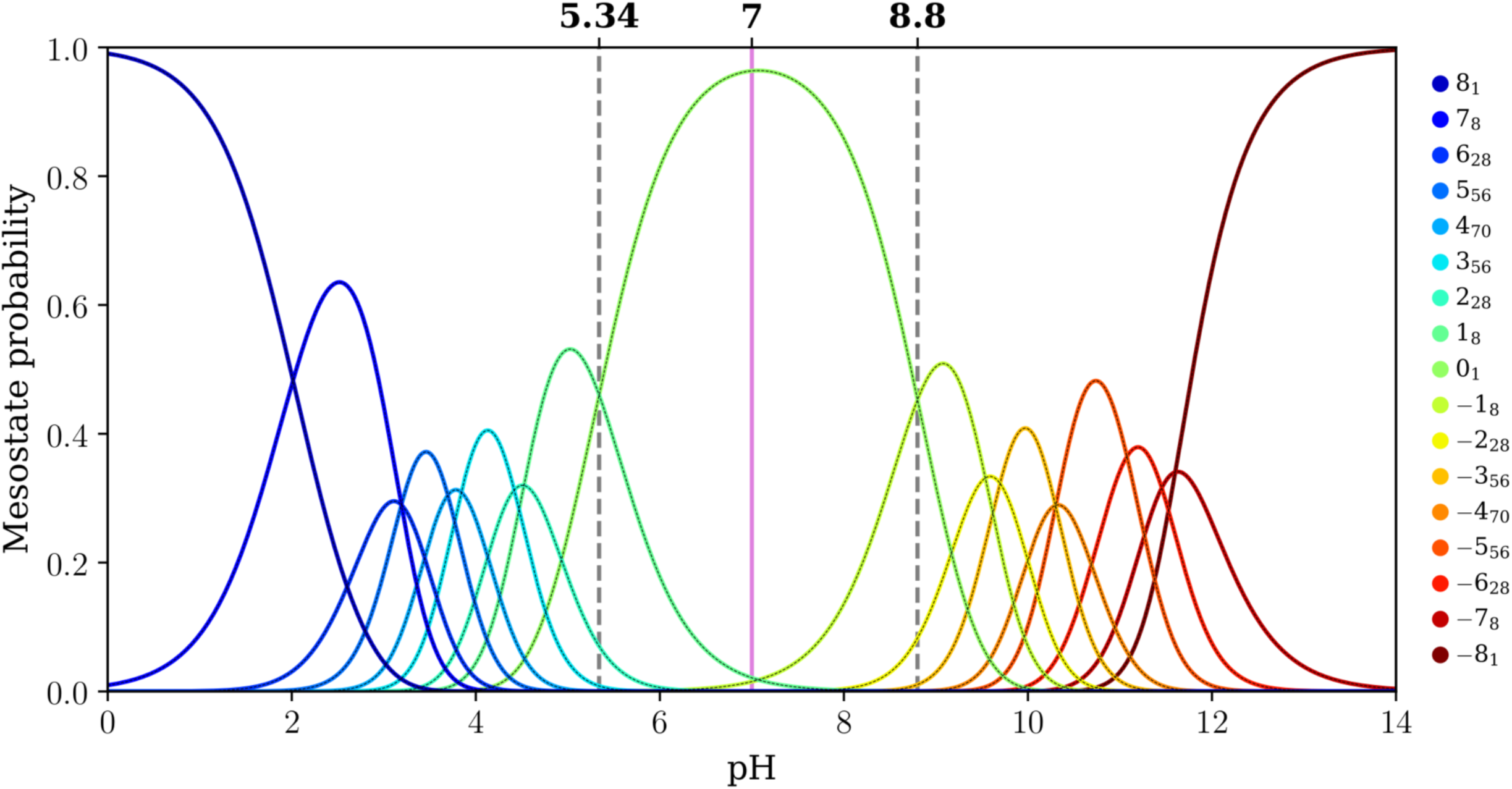
Mesostate probabilities, plotted against pH, for the E_4_K_4_ peptide. The dashed lines in each panel mark the pH values at which mesostates adjacent to the 0_1_ mesostate have equal likelihoods of being populated. The pink line in each panel corresponds to a pH of 7.0.

**Figure 7:**
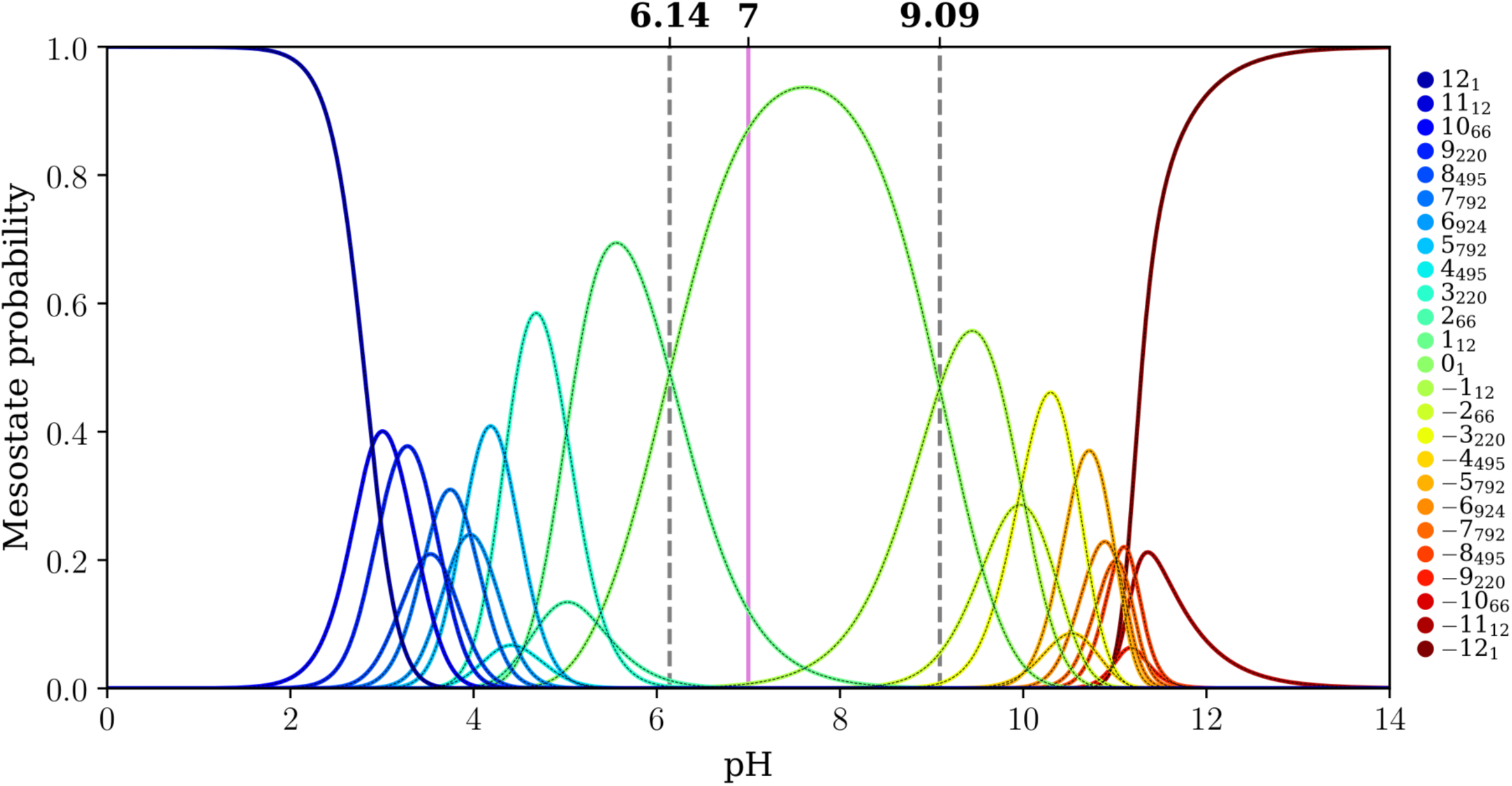
Mesostate probabilities, plotted against pH, for the (E_4_K_4_)_2_ peptide. The dashed lines in each panel mark the pH values at which mesostates adjacent to the 0_1_ mesostate have equal likelihoods of being populated. The pink line in each panel corresponds to a pH of 7.0.

**Figure 8:**
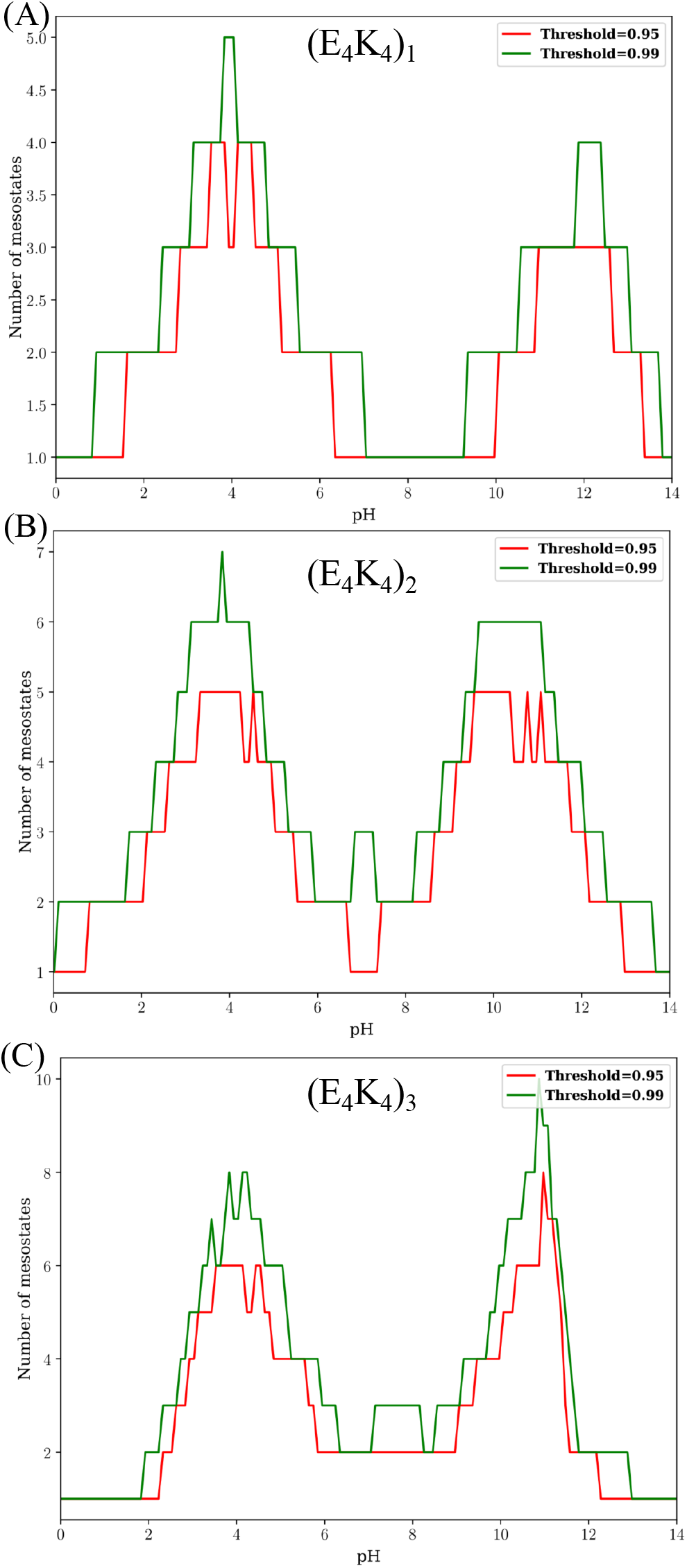
Mesostate probabilities, plotted against pH, for the (E_4_K_4_)_3_ peptide. The dashed lines in each panel mark the pH values at which mesostates adjacent to the 0_1_ mesostate have equal likelihoods of being populated. The pink line in each panel corresponds to a pH of 7.0.

For (E_4_K_4_)_2_ the pK_a_ values for the transitions +1_8_ ↔ 0_1_ and 0_1_ ↔ −1_8_ are 5.34 and 8.80, respectively (**Figure 7**) implying a gap of 3.46 pH units. This gap between the two pK_a_ values, which corresponds to transitions between 0_1_ and its adjacent mesostates, further narrows for (E_4_K_4_)_3_ (**Figure 8**). Here, the pK_a_ values for the transitions +1_12_ ↔ 0_1_ and 0_1_ ↔ −1_12_ are 6.14 and 9.01, respectively (gap of 2.87 pH units). The mesostate pK_a_ for transitions between the mesostate with net charge +1 and the 0_1_ mesostate shifts up with increasing numbers of repeats. In contrast, the pK_a_ for transitions between the mesostate with net charge −1 and 0_1_ shows non-monotonic variation with the number of repeats. This derives from the steeper variation of net charge with pH as the intrinsic pK_a_ of Lys is approached. In contrast, the mesostate with charge +1 is accessible in a pH range where the net charge shows a weaker pH dependence.

The gap between pK_a_ values for transitions between the 0_1_ and adjacent mesostates that have a net charge of ±1 narrows from 6.33 to 3.46 and 2.87 pH units as the number of E_4_K_4_repeats increases from 1 to 2 to 3. It is likely that this gap further narrows as the numbers of repeats increase. One of the main takeaways is that for (E_4_K_4_)_3_, the mesostate +1_12_ contributes significantly, in addition to the 0_1_, mesostate at pH 7.0 (see pink line in **Figure 8**). This contribution weakens in favor of the −1_12_ mesostate as the pH increases above 7.4. Therefore, even at pH 7.0, there are at least 13 distinct microstates whose contributions to the conformational ensemble must be considered.

The importance of charge state heterogeneity increases as the pH increases or decreases from the value of 7.0. The pH dependent populations of mesostates can be used to quantify the numbers of mesostates that contribute to 95% vs. 99% of the overall population as a function of pH. The results are summarized in **Figure 9**. **Figure 9A** shows that a single mesostate, 0_1_, and hence a single microstate, contributes to 99% of the population for the E_4_K_4_ peptide across a wide pH range. For the (E_4_K_4_)_2_, 2-3 mesostates are required to account for 90-99% of the preferred mesostates between the pH range of 7-9 (**Figure 9B**). We obtain similar results for the (E_4_K_4_)_3_ system (**Figure 9C**).

## DISCUSSION

Here, we have introduced a method for analyzing potentiometric data by leveraging the formal structure of *q*-canonical ensemble, which allows us to decouple analyses of measurements of charge from measurements of conformations. The *q*-canonical ensemble formally accounts for the contributions of charge state heterogeneity. Our methods are likely to be of particular use for uncovering the dual impacts of charge state and conformational heterogeneity for IDPs, as these systems tend to be rich in ionizable residues, with a large fraction featuring more than 30% of ionizable residues (19). This number increases when we include the contributions of neutral residues that become ionizable following post-translational modifications such as Ser / Thr phosphorylation.

The prothymosin α system that has been studied extensively by the Schuler group (27, 97-102) as well as acid-rich disordered proteins that function as transactivation domains of transcription factors (103), are examples of well-known IDPs that are among the closest mimics of homo polyacids, which are seldom fully deprotonated in solution (53, 54). Indeed, single molecule electrometry measurements performed by Ruggeri et al., (101) suggest that only 28.5 ± 1.2 out of the 46 D/E residues in prothymosin α are, on average, deprotonated at pH 8.8. Given the growing interest in the conformational, binding (104), and phase equilibria of polyampholytic, polyzwitterionic (105, 106), and polyelectrolytic IDPs (107), it will be important to measure net charge profiles and extract the contributions of charge state heterogeneity to these profiles. Here, we demonstrate the feasibility of such analyses by extracting mesostate populations using the *q*-canonical ensemble formalism. In follow up work, we propose to adopt this approach for analysis of cross-sections of IDPs.

The apparent insensitivity of helicity of so-called single alpha helices to changes in pH (94, 96) might derive from the helicity associated with microstates that have a net charge. In this scenario, it cannot solely be the network of salt bridges that contribute to stabilizing helical conformations. Instead, the interplay among differences in free energies of solvation of charged and neutral forms of Glu and Lys (108) as well as the higher intrinsic helical propensity of neutral Glu vs. charged Glu (109–112) are likely contributors to the intriguing robustness of measured helicities to large-scale changes in pH. Accounting for the totality of these interactions requires a combination of charge measurements, presented here, and measurements of conformational equilibria, that are jointly used in a suitable molecular simulation framework for describing the total *q*-canonical ensemble.

### Features of the *q*-canonical ensemble

The structure of the *q*-canonical ensemble allows us to decouple measurements of charge from measurements of conformation. By leveraging this decoupling, specifically the mesostate description of the net charge extracted from the *q*-canonical ensemble, we can analyze potentiometry-based charge profiles as a function of pH to quantify the number and types of mesostates that are thermodynamically relevant for a given set of solution conditions. We find that microstate entropy per mesostate makes significant contributions to the distribution of thermodynamically accessible mesostates. Importantly, the contributions of microstate entropy grow with the number of ionizable residues. Whether or not this heterogeneity can be overcome by dominant conformational preferences of specific microstates can only be decided by decoupling the measurements of charge from other readouts such as measurements of conformation and / or binding.

It is worth emphasizing the intrinsic differences between a site-specific representation of ionizable groups, and a description in terms of mesostates afforded by the *q*-canonical ensemble. If one were to measure pK_a_ values using site-specific titrations, aided by appropriate spectroscopic approaches such as nuclear magnetic resonance (NMR) (8, 81, 113) or infrared (IR) spectroscopy (111), the inferred variations in site-specific pK_a_ values would be much lower than the variations that we can uncover using the mesostate representation and measurements of global charge profiles.

### Potentiometry

The simplicity of potentiometric titrations is key to its robustness (114). The buffering capacity can be measured to obtain net charge profiles as a function of pH by fixing solution conditions such as salt concentration, salt type, and the solution temperature. Therefore, with suitable automation, potentiometry can provide direct access to charge profiles as a function of pH for a range of systems and solution conditions. These data, when analyzed using the *q*-canonical formalism, will be essential for understanding the extent of sequence- and composition-specific effects of charge regulation via proton binding and release.

Being one of the earliest methods to characterize acid-base equilibria in chemical species, several methods for the interpretation of potentiometric curves, especially for monoacids, have been developed (114). However, as noted by Ghasemi and Larson, the interpretation of potentiometric data has remained an unsolved problem for polyacids (54). We propose that our *q*-canonical ensemble-based approach provides an alternative route to obtain more complete descriptions of the measured charge profiles.

Historically, potentiometry measurements have been used to assess the presence of anomalous or shifted pK_a_ values (2). This is typically achieved by using model compound pK_a_ values, ignoring the microstate entropy, and querying whether the measured net charge profiles match or deviate from the calculated profiles. The qualitative assessments from analysis of potentiometry are typically used to set up detailed site-specific measurements of local pK_a_ values using NMR spectroscopy or other types of spectroscopies that rely on isotopic labeling at specific sites.

The net charge profiles extracted using potentiometry have also been analyzed by prescribing model conformations and the calculation of conformation-specific electrostatic potentials. The goal in these endeavors is to assess the extent to which local sequence contexts can be implicated as modifiers of local pK_a_ values that cause deviations of measured charge profiles from expectations based on model compounds. While these efforts highlight the importance of charge regulation, they ignore the contributions of microstate entropy. Here, we leverage the structure of the *q*-canonical ensemble and show how this can be deployed to extract insights regarding the pH-dependence of mesostate populations.

### Potentiometry vs. other methods to measure charge

Methods to measure charge are of considerable interest, especially for the study of IDPs. Several methods such as capillary electrophoresis (57, 115) and single molecule electrometry (101) have been introduced and deployed for measuring the net charges of IDPs and other flexible polymers. A limitation of these methods is the reliance on measurements of molecular mobility in the presence of a potential drop or a spatially patterned electric field. Analysis of data from mobility-based measurements requires the *a priori* assumption of a preferred conformational state. Further complications arise because electrophoretic mobility measurements require the use of ultra-low salt concentrations to avoid confounding effects that arise from the adsorption or release of solution ions. Accordingly, we propose that potentiometry, which is the most direct approach for measuring net charge, should be the preferred route for quantifying charge profiles of IDPs.

### Summary and prognosis

We have developed and deployed a formalism, leveraging the recently introduced *q*-canonical ensemble, to analyze potentiometric titrations. We showcase the approach using measurements and analyses for three related systems. The formalism is applicable for analyzing multisite binding isotherms (116–118) for arbitrary ligands, not just protons. A large-scale deployment of potentiometry, aided by the analysis introduced in this work, will be forthcoming for a cross-section of IDPs that include residues such as Asp, Glu, Lys, Arg, His, and other more complex features. This will allow us to dissect the contributions of different compositions and sequence contexts to charge state heterogeneity.

The results presented here help highlight the importance of charge state heterogeneity and the fact that heterogeneity makes significant contributions with increasing numbers of ionizable residues. Potentiometric measurements, combined with separate measurements of the pH dependence of conformational properties, can be analyzed using the *q*-canonical ensemble to extract mesostate populations as well as microstate populations. The latter is being pursued for a series of systems, using sampling methods developed for the *q*-canonical ensemble (70). The results, which will be published elsewhere, will likely move us in the direction of quantifying the extent to which charge state and conformational heterogeneity either enhance or antagonize one another. These studies are likely to be important for understanding and modeling the conformational, binding, and phase equilibria of IDPs. Such studies take on additional significance given various observations of pH-responsive equilibria for IDPs (42), recent measurements highlighting the significant extent of charge regulation in polyelectrolytic IDPs, and the presence of compartments of highly variable pH environments within cells.

## AUTHOR CONTRIBUTIONS

Conceptualization: M.J.F., and R.V.P. Development of the formal theory and analysis: M.J.F. Potentiometry setup, troubleshooting, and measurements: A.E.P. Numerical analysis of potentiometry data: M.J.F., and R.V.P. Writing of the manuscript: M.J.F., and R.V.P. Editing: M.J.F., A.E.P., and R.V.P. Funding: R.V.P.

## ACKNOWLEDGMENTS

We thank Furqan Dar, Mina Farag, Andrew Lin, Kiersten Ruff, Min Kyung Shinn, and Xiangze Zeng for careful reading of the manuscript. This work was supported by grant FA9550-20-1-0241 from the Air Force Office of Scientific Research, grants R01NS089932 and 5R01NS056114 from the US National Institutes of Health, and the St. Jude Children’s Research Collaborative on Membraneless Organelles.

---

**Figure 9: Minimum number of mesostates that are required to account for 95% and 99% of the population, as a function of pH, for each of the three peptides.**

## Notes

### Competing Interest Statement

The authors have declared no competing interest.

### Summary of Updates

This version of the manuscript has been revised in response to feedback from reviewers.

## REFERENCES

1. Pace, C. N., G. R. Grimsley, and J. M. Scholtz. 2009. Protein Ionizable Groups: pK Values and Their Contribution to Protein Stability and Solubility. Journal of Biological Chemistry 284:13285–13289.

2. Whitten, S. T., and E. B. Garcia-Moreno. 2000. pH dependence of stability of staphylococcal nuclease: evidence of substantial electrostatic interactions in the denatured state. Biochemistry 39:14292–14304.

3. Whitten, S. T., J. O. Wooll, R. Razeghifard, E. B. Garcia-Moreno, and V. J. Hilser. 2001. The origin of pH-dependent changes in m-values for the denaturant-induced unfolding of proteins. Journal of molecular biology 309:1165–1175.

4. Castle, P. E., D. A. Karp, L. Zeitlin, E. B. Garcia-Moreno, T. R. Moench, K. J. Whaley, and R. A. Cone. 2002. Human monoclonal antibody stability and activity at vaginal pH. Journal of reproductive immunology 56:61–76.

5. Mehler, E. L., M. Fuxreiter, I. Simon, and E. B. Garcia-Moreno. 2002. The role of hydrophobic microenvironments in modulating pKa shifts in proteins. Proteins: Structure, Function, Bioinformatics 48:283–292.

6. Schlessman, J. L., C. Abe, A. Gittis, D. A. Karp, M. A. Dolan, and E. B. Garcia-Moreno. 2008. Crystallographic study of hydration of an internal cavity in engineered proteins with buried polar or ionizable groups. Biophys J 94:3208–3216.

7. Whitten, S. T., E. B. Garcia-Moreno, and V. J. Hilser. 2005. Local conformational fluctuations can modulate the coupling between proton binding and global structural transitions in proteins. Proc Natl Acad Sci U S A 102:4282–4287.

8. Castaneda, C. A., C. A. Fitch, A. Majumdar, V. Khangulov, J. L. Schlessman, and B. E. Garcia-Moreno. 2009. Molecular determinants of the pKa values of Asp and Glu residues in staphylococcal nuclease. Proteins 77:570–588.

9. Harms, M. J., C. A. Castaneda, J. L. Schlessman, G. R. Sue, D. G. Isom, B. R. Cannon, and E. B. Garcia-Moreno. 2009. The pK(a) values of acidic and basic residues buried at the same internal location in a protein are governed by different factors. Journal of molecular biology 389:34–47.

10. Isom, D. G., C. A. Castaneda, B. R. Cannon, P. D. Velu, and E. B. Garcia-Moreno. 2010. Charges in the hydrophobic interior of proteins. Proceedings of the National Academy of Sciences of the United States of America USA 107:16096–16100.

11. Kitahara, R., K. Hata, A. Maeno, K. Akasaka, M. S. Chimenti, E. B. Garcia-Moreno, M. A. Schroer, C. Jeworrek, M. Tolan, R. Winter, J. Roche, C. Roumestand, K. Montet de Guillen, and C. A. Royer. 2011. Structural plasticity of staphylococcal nuclease probed by perturbation with pressure and pH. Proteins 79:1293–1305.

12. Peck, M. T., G. Ortega, J. N. De Luca-Johnson, J. L. Schlessman, A. C. Robinson, and E. B. Garcia-Moreno. 2017. Local Backbone Flexibility as a Determinant of the Apparent pKa Values of Buried Ionizable Groups in Proteins. Biochemistry 56:5338–5346.

13. Petrosian, S. A., and G. I. Makhatadze. 2000. Contribution of proton linkage to the thermodynamic stability of the major cold-shock protein of Escherichia coli CspA. Protein science: a publication of the Protein Society 9:387–394.

14. Permyakov, S. E., G. I. Makhatadze, R. Owenius, V. N. Uversky, C. L. Brooks, E. A. Permyakov, and L.J. Berliner. 2005. How to improve nature: study of the electrostatic properties of the surface of alpha-lactalbumin. Protein Engineering Design and Selection 18:425–433.

15. Chan, C. H., C. C. Wilbanks, G. I. Makhatadze, and K. B. Wong. 2012. Electrostatic contribution of surface charge residues to the stability of a thermophilic protein: benchmarking experimental and predicted pKa values. PloS one 7:e30296.

16. Hofer, F., J. Kraml, U. Kahler, A. S. Kamenik, and K. R. Liedl. 2020. Catalytic Site pKa Values of Aspartic, Cysteine, and Serine Proteases: Constant pH MD Simulations. Journal of chemical information and modeling 60:3030–3042.

17. Bartlett, G. J., C. T. Porter, N. Borkakoti, and J. M. Thornton. 2002. Analysis of Catalytic Residues in Enzyme Active Sites. Journal of molecular biology 324:105–121.

18. Harris, T. K., and G. J. Turner. 2002. Structural Basis of Perturbed pKa Values of Catalytic Groups in Enzyme Active Sites. IUBMB Life 53:85–98.

19. Holehouse, A. S., R. K. Das, J. N. Ahad, M. O. G. Richardson, and R. V. Pappu. 2017. CIDER: Resources to Analyze Sequence-Ensemble Relationships of Intrinsically Disordered Proteins. Biophysical Journal 112:16–21.

20. Pujato, M., C. Bracken, R. Mancusso, M. Cataldi, and M. L. Tasayco. 2005. pH Dependence of Amide Chemical Shifts in Natively Disordered Polypeptides Detects Medium-Range Interactions with Ionizable Residues. Biophysical Journal 89:3293–3302.

21. Mao, A. H., S. L. Crick, A. Vitalis, C. L. Chicoine, and R. V. Pappu. 2010. Net charge per residue modulates conformational ensembles of intrinsically disordered proteins. Proceedings of the National Academy of Sciences of the United States of America 107:8183–8188.

22. Mao, A. H., N. Lyle, and R. V. Pappu. 2013. Describing sequence-ensemble relationships for intrinsically disordered proteins. Biochemical Journal 449:307–318.

23. van der Lee, R., M. Buljan, B. Lang, R. J. Weatheritt, G. W. Daughdrill, A. K. Dunker, M. Fuxreiter, J. Gough, J. Gsponer, D. T. Jones, P. M. Kim, R. W. Kriwacki, C. J. Oldfield, R. V. Pappu, P. Tompa, V. N. Uversky, P. E. Wright, and M. M. Babu. 2014. Classification of Intrinsically Disordered Regions and Proteins. Chemical Reviews 114:6589–6631.

24. Marsh, J. A., B. Dancheck, M. J. Ragusa, M. Allaire, J. D. Forman-Kay, and W. Peti. 2010. Structural Diversity in Free and Bound States of Intrinsically Disordered Protein Phosphatase 1 Regulators. Structure 18:1094–1103.

25. Marsh, J. A., and J. D. Forman-Kay. 2010. Sequence Determinants of Compaction in Intrinsically Disordered Proteins. Biophysical Journal 98:2383–2390.

26. Liu, B., D. Chia, V. Csizmok, P. Farber, J. D. Forman-Kay, and C. C. Gradinaru. 2014. The Effect of Intrachain Electrostatic Repulsion on Conformational Disorder and Dynamics of the Sic1 Protein. Journal of Physical Chemistry B 118:4088–4097.

27. Borgia, A., M. B. Borgia, K. Bugge, V. M. Kissling, P. O. Heidarsson, C. B. Fernandes, A. Sottini, A. Soranno, K. J. Buholzer, D. Nettels, B. B. Kragelund, R. B. Best, and B. Schuler. 2018. Extreme disorder in an ultrahigh-affinity protein complex. Nature 555:61–66.

28. Bugge, K., I. Brakti, C. B. Fernandes, J. E. Dreier, J. E. Lundsgaard, J. G. Olsen, K. Skriver, and B. B. Kragelund. 2020. Interactions by Disorder – A Matter of Context. Frontiers in Molecular Biosciences 7.

29. O’Shea, C., L. Staby, S. K. Bendsen, F. G. Tidemand, A. Redsted, M. Willemoës, B. B. Kragelund, and K. Skriver. 2017. Structures and Short Linear Motif of Disordered Transcription Factor Regions Provide Clues to the Interactome of the Cellular Hub Protein Radical-induced Cell Death1. Journal of Biological Chemistry 292:512–527.

30. Sottini, A., A. Borgia, M. B. Borgia, K. Bugge, D. Nettels, A. Chowdhury, P. O. Heidarsson, F. Zosel, R. B. Best, B. B. Kragelund, and B. Schuler. 2020. Polyelectrolyte interactions enable rapid association and dissociation in high-affinity disordered protein complexes. Nature communications 11:5736.

31. Prestel, A., N. Wichmann, J. M. Martins, R. Marabini, N. Kassem, S. S. Broendum, M. Otterlei, O. Nielsen, M. Willemoës, M. Ploug, W. Boomsma, and B. B. Kragelund. 2019. The PCNA interaction motifs revisited: thinking outside the PIP-box. Cellular and Molecular Life Sciences 76:4923–4943.

32. Millard, P. S., K. Weber, B. B. Kragelund, and M. Burow. 2019. Specificity of MYB interactions relies on motifs in ordered and disordered contexts. Nucleic Acids Research 47:9592–9608.

33. Borg, M., T. Mittag, T. Pawson, M. Tyers, J. D. Forman-Kay, and H. S. Chan. 2007. Polyelectrostatic interactions of disordered ligands suggest a physical basis for ultrasensitivity. Proceedings of the National Academy of Sciences of the United States of America 104:9650–9655.

34. Pak, C. W., M. Kosno, A. S. Holehouse, S. B. Padrick, A. Mittal, R. Ali, A. A. Yunus, D. R. Liu, R. V. Pappu, and M. K. Rosen. 2016. Sequence Determinants of Intracellular Phase Separation by Complex Coacervation of a Disordered Protein. Molecular cell 63:72–85.

35. Kaur, T., M. Raju, I. Alshareedah, R. B. Davis, D. A. Potoyan, and P. R. Banerjee. 2021. Sequence-encoded and composition-dependent protein-RNA interactions control multiphasic condensate morphologies. Nature communications 12:872.

36. Alshareedah, I., M. M. Moosa, M. Raju, D. A. Potoyan, and P. R. Banerjee. 2020. Phase transition of RNA-protein complexes into ordered hollow condensates. Proceedings of the National Academy of Sciences 117:15650.

37. Fisher, R. S., and S. Elbaum-Garfinkle. 2020. Tunable multiphase dynamics of arginine and lysine liquid condensates. Nature communications 11:4628.

38. Yeong, V., J.-w. Wang, J. M. Horn, and A. C. Obermeyer. 2021. Intracellular phase separation of globular proteins facilitated by short cationic peptides. bioRxiv:2021.2007.2008.450573.

39. Chang, L. W., T. K. Lytle, M. Radhakrishna, J. J. Madinya, J. Velez, C. E. Sing, and S. L. Perry. 2017. Sequence and entropy-based control of complex coacervates. Nature communications 8:1273.

40. Knoerdel, A. R., W. C. Blocher McTigue, and C. E. Sing. 2021. Transfer Matrix Model of pH Effects in Polymeric Complex Coacervation. The Journal of Physical Chemistry B 125:8965–8980.

41. Last, M. G. F., S. Deshpande, and C. Dekker. 2020. pH-Controlled Coacervate–Membrane Interactions within Liposomes. ACS Nano 14:4487–4498.

42. Quiroz, F. G., V. F. Fiore, J. Levorse, L. Polak, E. Wong, H. A. Pasolli, and E. Fuchs. 2020. Liquid-liquid phase separation drives skin barrier formation. Science (New York, N.Y.) 367:eaax9554.

43. Lund, M., and B. Jönsson. 2013. Charge regulation in biomolecular solution. Quarterly Reviews of Biophysics 46:265–281.

44. Avni, Y., D. Andelman, and R. Podgornik. 2019. Charge regulation with fixed and mobile charged macromolecules. Current Opinion in Electrochemistry 13:70–77.

45. Burton, R. F. 1988. The protein content of extracellular fluids and its relevance to the study of ionic regulation: Net charge and colloid osmotic pressure. Comparative Biochemistry and Physiology Part A: Physiology 90:11–16.

46. Bah, A., and J. D. Forman-Kay. 2016. Modulation of Intrinsically Disordered Protein Function by Post-translational Modifications. Journal of Biological Chemistry 291:6696–6705.

47. Gil, J., A. Ramírez-Torres, and S. Encarnación-Guevara. 2017. Lysine acetylation and cancer: A proteomics perspective. Journal of Proteomics 150:297–309.

48. Fuhrmann, J., and P. R. Thompson. 2016. Protein Arginine Methylation and Citrullination in Epigenetic Regulation. ACS chemical biology 11:654–668.

49. Bah, A., R. M. Vernon, Z. Siddiqui, M. Krzeminski, R. Muhandiram, C. Zhao, N. Sonenberg, L. E. Kay, and J. D. Forman-Kay. 2015. Folding of an intrinsically disordered protein by phosphorylation as a regulatory switch. Nature 519:106–109.

50. Kennelly, P. J. 2014. Protein Ser/Thr/Tyr Phosphorylation in the Archaea*. Journal of Biological Chemistry 289:9480–9487.

51. Eymann, C., D. Becher, J. Bernhardt, K. Gronau, A. Klutzny, and M. Hecker. 2007. Dynamics of protein phosphorylation on Ser/Thr/Tyr in Bacillus subtilis. PROTEOMICS 7:3509–3526.

52. Bischoff, R., and H. V. J. Kolbe. 1994. Deamidation of asparagine and glutamine residues in proteins and peptides: structural determinants and analytical methodology. Journal of Chromatography B: Biomedical Sciences and Applications 662:261–278.

53. Salehi, A., and R. G. Larson. 2016. A Molecular Thermodynamic Model of Complexation in Mixtures of Oppositely Charged Polyelectrolytes with Explicit Account of Charge Association/Dissociation. Macromolecules 49:9706–9719.

54. Ghasemi, M., and R. G. Larson. 2021. Role of electrostatic interactions in charge regulation of weakly dissociating polyacids. Progress in Polymer Science 112:101322.

55. Fitch, C. A., S. T. Whitten, V. J. Hilser, and E. B. Garcia-Moreno. 2006. Molecular mechanisms of pH-driven conformational transitions of proteins: insights from continuum electrostatics calculations of acid unfolding. Proteins 63:113–126.

56. Hilser, V. J., E. B. Garcia-Moreno, T. G. Oas, G. Kapp, and S. T. Whitten. 2006. A statistical thermodynamic model of the protein ensemble. Chemical Reviews 106:1545–1558.

57. Sharma, U., R. S. Negin, and J. D. Carbeck. 2003. Effects of Cooperativity in Proton Binding on the Net Charge of Proteins in Charge Ladders. The Journal of Physical Chemistry B 107:4653–4666.

58. Simonson, T., J. Carlsson, and D. A. Case. 2004. Proton Binding to Proteins: pKa Calculations with Explicit and Implicit Solvent Models. Journal of the American Chemical Society 126:4167–4180.

59. Shiao, D. D. F., and J. M. Sturtevant. 1976. Heats of binding protons to globular proteins. Biopolymers 15:1201–1211.

60. Anderson, C. F., and M. T. Record. 1993. Salt dependence of oligoion-polyion binding: a thermodynamic description based on preferential interaction coefficients. The Journal of Physical Chemistry 97:7116–7126.

61. Grosberg, A. Y., T. T. Nguyen, and B. I. Shklovskii. 2002. Colloquium: The physics of charge inversion in chemical and biological systems. Reviews of Modern Physics 74:329–345.

62. Roosen-Runge, F., B. S. Heck, F. Zhang, O. Kohlbacher, and F. Schreiber. 2013. Interplay of pH and Binding of Multivalent Metal Ions: Charge Inversion and Reentrant Condensation in Protein Solutions. The Journal of Physical Chemistry B 117:5777–5787.

63. Zhang, F., S. Weggler, M. J. Ziller, L. Ianeselli, B. S. Heck, A. Hildebrandt, O. Kohlbacher, M. W. A. Skoda, R. M. J. Jacobs, and F. Schreiber. 2010. Universality of protein reentrant condensation in solution induced by multivalent metal ions. Proteins: Structure, Function, and Bioinformatics 78:3450–3457.

64. Winkler, R. G., M. Gold, and P. Reineker. 1998. Collapse of Polyelectrolyte Macromolecules by Counterion Condensation and Ion Pair Formation: A Molecular Dynamics Simulation Study. Physical review letters 80:3731–3734.

65. Butler, J. C., T. Angelini, J. X. Tang, and G. C. L. Wong. 2003. Ion Multivalence and Like-Charge Polyelectrolyte Attraction. Physical review letters 91:028301.

66. Angelini, T. E., H. Liang, W. Wriggers, and G. C. L. Wong. 2003. Like-charge attraction between polyelectrolytes induced by counterion charge density waves. Proceedings of the National Academy of Sciences 100:8634.

67. Annunziata, O., L. Paduano, and J. G. Albright. 2007. Multicomponent Diffusion of Lysozyme in Aqueous Calcium Chloride. The Role of Common-Ion Effects and Protein-Salt Preferential Interactions. The Journal of Physical Chemistry B 111:10591–10598.

68. Zheng, Y., and Q. Cui. 2017. Microscopic mechanisms that govern the titration response and pKa values of buried residues in staphylococcal nuclease mutants. Proteins: Structure, Function, and Bioinformatics 85:268–281.

69. Pahari, S., L. Sun, and E. Alexov. 2019. PKAD: a database of experimentally measured pKa values of ionizable groups in proteins. Database 2019.

70. Fossat, M. J., and R. V. Pappu. 2019. q-Canonical Monte Carlo Sampling for Modeling the Linkage between Charge Regulation and Conformational Equilibria of Peptides. The Journal of Physical Chemistry B 123:6952–6967.

71. Mandel, M., and J. C. Leyte. 1972. Some remarks on the titration equation of weak polyacids. Journal of Electroanalytical Chemistry and Interfacial Electrochemistry 37:297–301.

72. Müller, M., L. Wirth, and B. Urban. 2021. Determination of the Carboxyl Dissociation Degree and pKa Value of Mono and Polyacid Solutions by FTIR Titration. Macromolecular Chemistry and Physics 222:2000334.

73. Borukhov, I., D. Andelman, R. Borrega, M. Cloitre, L. Leibler, and H. Orland. 2000. Polyelectrolyte Titration: Theory and Experiment. The Journal of Physical Chemistry B 104:11027–11034.

74. Woodbury, C. P. 1993. The titration curve of weak polyacids. The Journal of Physical Chemistry 97:3623–3630.

75. Fitch, C. A., G. Platzer, M. Okon, B. Garcia-Moreno E., and L. P. McIntosh. 2015. Arginine: Its pKa value revisited. Protein science: a publication of the Protein Society 24:752–761.

76. Bakker, E., and E. Pretsch. 2007. Modern Potentiometry. Angewandte Chemie International Edition 46:5660–5668.

77. Smith, L. A., M. W. Glasscott, K. J. Vannoy, and J. E. Dick. 2020. Enzyme Kinetics via Open Circuit Potentiometry. Analytical Chemistry 92:2266–2273.

78. Gunner, M. R., T. Murakami, A. S. Rustenburg, M. Isik, and J. D. Chodera. 2020. Standard state free energies, not pKas, are ideal for describing small molecule protonation and tautomeric states. Journal of Computer-Aided Molecular Design 34:561–573.

79. Szakács, Z., M. Kraszni, and B. Noszál. 2004. Determination of microscopic acid–base parameters from NMR–pH titrations. Analytical and Bioanalytical Chemistry 378:1428–1448.

80. Bashford, D., and M. Karplus. 1991. Multiple-site titration curves of proteins: an analysis of exact and approximate methods for their calculation. The Journal of Physical Chemistry 95:9556–9561.

81. Onufriev, A., D. A. Case, and G. M. Ullmann. 2001. A Novel View of pH Titration in Biomolecules. Biochemistry 40:3413–3419.

82. Metropolis, N., A. W. Rosenbluth, M. N. Rosenbluth, A. H. Teller, and E. Teller. 1953. Equation of State Calculations by Fast Computing Machines. The Journal of Chemical Physics 21:1087–1092.

83. Platzer, G., M. Okon, and L. P. McIntosh. 2014. pH-dependent random coil 1H, 13C, and 15N chemical shifts of the ionizable amino acids: a guide for protein pKa measurements. Journal of Biomolecular NMR 60:109–129.

84. Savitzky, A., and M. J. E. Golay. 1964. Smoothing and Differentiation of Data by Simplified Least Squares Procedures. Analytical Chemistry 36:1627–1639.

85. Gran, G. 1952. Determination of the equivalence point in potentiometric titrations. Part II. Analyst 77:661–671.

86. Gran, G., and A. Johansson. 1981. Automatic titration by stepwise addition of equal volumes of titrant. Part VI. Further extension of the Gran I method for calculation of the equivalence volume in acid-base titrations. Analyst 106:231–242.

87. Yan, J. F. 1965. A Method for the Determination of Equivalence Point in Potentiometric Titrations Using Unequal Volume Increments. Analytical Chemistry 37:1588–1590.

88. Fortuin, J. M. H. 1961. Method for determination of the equivalence point in potentiometric titrations. Analytica Chimica Acta 24:175–191.

89. Nozaki, Y., and C. Tanford. 1967. Examination of titration behavior. In Methods in enzymology. Academic Press. 715–734.

90. Huang, Y., and D. W. Bolen. 1995. Probes of energy transduction in enzyme catalysis. In Methods in enzymology. Academic Press. 19–43.

91. Baker, E. G., G. J. Bartlett, M. P. Crump, R. B. Sessions, N. Linden, C. F. J. Faul, and D. N. Woolfson. 2015. Local and macroscopic electrostatic interactions in single α-helices. Nature Chemical Biology 11:221–228.

92. Peckham, M., and P. J. Knight. 2009. When a predicted coiled coil is really a single a-helix, in myosins and other proteins. Soft Matter 5:2493–2503.

93. Süveges, D., Z. Gáspári, G. Tóth, and L. Nyitray. 2009. Charged single α-helix: A versatile protein structural motif. Proteins: Structure, Function, and Bioinformatics 74:905–916.

94. Wolny, M., M. Batchelor, G. J. Bartlett, E. G. Baker, M. Kurzawa, P. J. Knight, L. Dougan, D. N. Woolfson, E. Paci, and M. Peckham. 2017. Characterization of long and stable de novo single alpha-helix domains provides novel insight into their stability. Scientific Reports 7:44341.

95. Malik, R. U., M. Dysthe, M. Ritt, R. K. Sunahara, and S. Sivaramakrishnan. 2017. ER/K linked GPCR-G protein fusions systematically modulate second messenger response in cells. Scientific Reports 7:7749.

96. Wang, E., and C.-L. A. Wang. 1996. (i,i+ 4) Ion Pairs Stabilize Helical Peptides Derived from Smooth Muscle Caldesmon. Archives of Biochemistry and Biophysics 329:156–162.

97. Mueller-Spaeth, S., A. Soranno, V. Hirschfeld, H. Hofmann, S. Rueegger, L. Reymond, D. Nettels, and B. Schuler. 2010. Charge interactions can dominate the dimensions of intrinsically disordered proteins. Proceedings of the National Academy of Sciences of the United States of America 107:14609–14614.

98. Hofmann, H., A. Soranno, A. Borgia, K. Gast, D. Nettels, and B. Schuler. 2012. Polymer scaling laws of unfolded and intrinsically disordered proteins quantified with singlemolecule spectroscopy. Proceedings of the National Academy of Sciences of the United States of America 109:16155–16160.

99. Brucale, M., B. Schuler, and B. Samori. 2014. Single-Molecule Studies of Intrinsically Disordered Proteins. Chemical Reviews 114:3281–3317.

100. Wuttke, R., H. Hofmann, D. Nettels, M. B. Borgia, J. Mittal, R. B. Best, and B. Schuler. 2014. Temperature-dependent solvation modulates the dimensions of disordered proteins. Proceedings of the National Academy of Sciences of the United States of America 111:5213–5218.

101. Ruggeri, F., F. Zosel, N. Mutter, M. Rozycka, M. Wojtas, A. Ozyhar, B. Schuler, and M. Krishnan. 2017. Single-molecule electrometry. Nature Nanotechnology 12:488–495.

102. Holmstrom, E. D., A. Holla, W. Zheng, D. Nettels, R. B. Best, and B. Schuler. 2018. Accurate Transfer Efficiencies, Distance Distributions, and Ensembles of Unfolded and Intrinsically Disordered Proteins From Single-Molecule FRET. Methods in enzymology 611:287–325.

103. Staller, M. V., A. S. Holehouse, D. Swain-Lenz, R. K. Das, R. V. Pappu, and B. A. Cohen. 2018. A High-Throughput Mutational Scan of an Intrinsically Disordered Acidic Transcriptional Activation Domain. Cell systems 6:444–455.e446.

104. Dyla, M., and M. Kjaergaard. 2020. Intrinsically disordered linkers control tethered kinases via effective concentration. Proceedings of the National Academy of Sciences 117:21413.

105. Sørensen, C. S., and M. Kjaergaard. 2019. Effective concentrations enforced by intrinsically disordered linkers are governed by polymer physics. Proceedings of the National Academy of Sciences 116:23124.

106. Greig, J. A., T. A. Nguyen, M. Lee, A. S. Holehouse, A. E. Posey, R. V. Pappu, and G. Jedd. 2020. Arginine-Enriched Mixed-Charge Domains Provide Cohesion for Nuclear Speckle Condensation. Molecular cell 77:1237–1250.e1234.

107. Wiggers, F., S. Wohl, A. Dubovetskyi, G. Rosenblum, W. Zheng, and H. Hofmann. 2021. Diffusion of the disordered E-cadherin tail on β–catenin. bioRxiv:2021.2002.2003.429507.

108. Fossat, M. J., X. Zeng, and R. V. Pappu. 2021. Uncovering Differences in Hydration Free Energies and Structures for Model Compound Mimics of Charged Side Chains of Amino Acids. The Journal of Physical Chemistry B 125:4148–4161.

109. Nick Pace, C., and J. Martin Scholtz. 1998. A Helix Propensity Scale Based on Experimental Studies of Peptides and Proteins. Biophysical Journal 75:422–427.

110. Myer, Y. P. 1969. The pH–Induced Helix–Coil Transition of Poly–L–lysine and Poly–L–glutamic Acid and the 238–mµ Dichroic Band. Macromolecules 2:624–628.

111. Donten, M. L., and P. Hamm. 2013. pH-jump induced α-helix folding of poly-l-glutamic acid. Chemical Physics 422:124–130.

112. Fonin, A. V., O. V. Stepanenko, A. K. Sitdikova, I. A. Antifeeva, E. I. Kostyleva, A. M. Polyanichko, M. M. Karasev, S. A. Silonov, O. I. Povarova, I. M. Kuznetsova, V. N. Uversky, and K. K. Turoverov. 2019. Folding of poly-amino acids and intrinsically disordered proteins in overcrowded milieu induced by pH change. International Journal of Biological Macromolecules 125:244–255.

113. Donnini, S., R. T. Ullmann, G. Groenhof, and H. Grubmuller. 2016. Charge-Neutral Constant pH Molecular Dynamics Simulations Using a Parsimonious Proton Buffer. J Chem Theory Comput 12:1040–1051.

114. Buck, R. P. 1972. Ion selective electrodes, potentiometry, and potentiometric titrations. Analytical Chemistry 44:270–295.

115. Plasson, R., and H. Cottet. 2006. Determination and Modeling of Peptide pKa by Capillary Zone Electrophoresis. Analytical Chemistry 78:5394–5402.

116. Gill, S. J., B. Richey, G. Bishop, and J. Wyman. 1985. Generalized binding phenomena in an allosteric macromolecule. Biophysical Chemistry 21:1–14.

117. Di Cera, E., S. J. Gill, and J. Wyman. 1988. Binding capacity: cooperativity and buffering in biopolymers. Proceedings of the National Academy of Sciences 85:449.

118. Gill, S. J., H. T. Gaud, J. Wyman, and B. G. Barisas. 1978. Analysis of ligand binding curves in terms of species fractions. Biophysical Chemistry 8:53–59.

