## Supporting Information for "Quantifying charge state heterogeneity for proteins with multiple ionizable residues"

**for**

**Movie S1: Movie depicting the evolution of the fit as a function of the number of fitting steps for all $pK_a$ values starting at 7.** The grey curve is the smoothed, interpolated experimental data, the orange curve is the current fit. The red dashed lines correspond to the fitted $pK_a$ values which in the q-canonical approach are defined as the pH at which two mesostates with a net charge difference of one have an equal population.

**Movie S2: Movie depicting the evolution of the fit as a function of the number of fitting steps for all $pK_a$ values starting from intrinsic $pK_a$ values.** Colors are the same as in Movie S1.

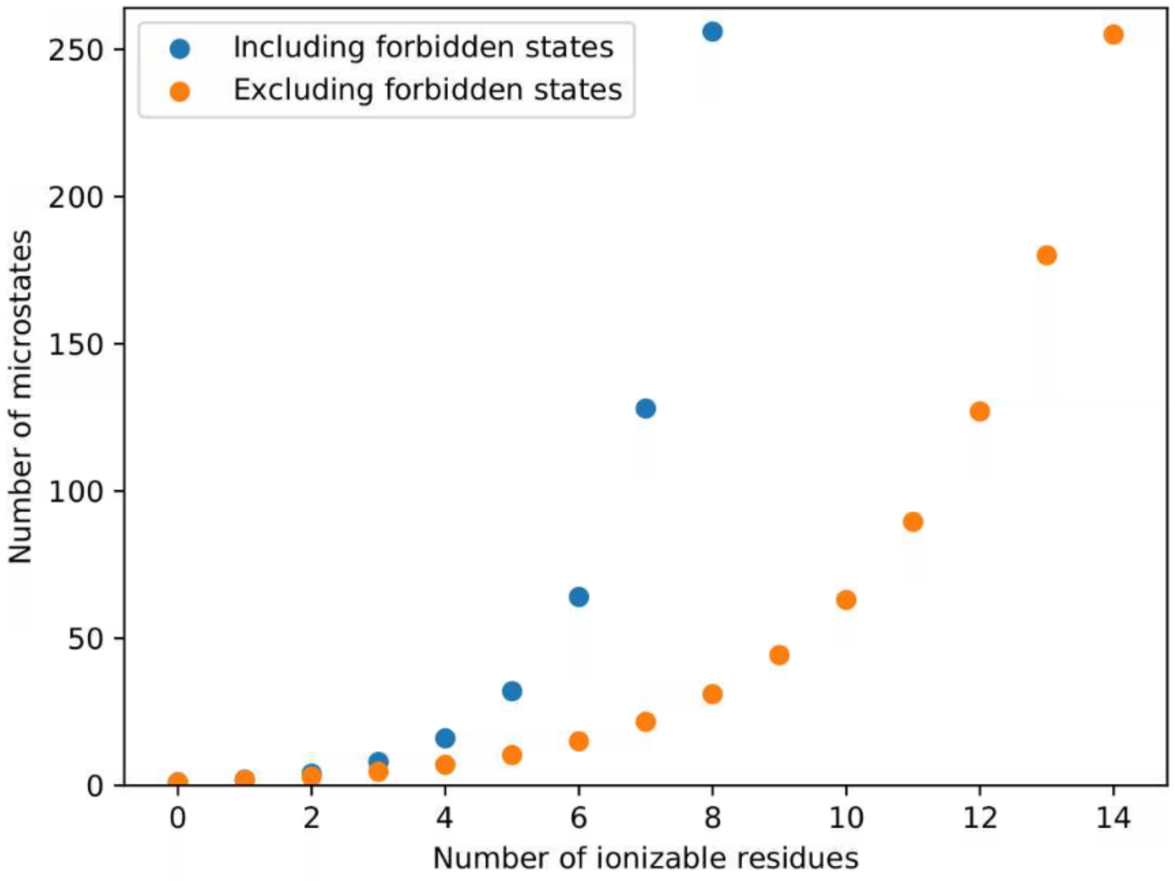


**Figure S1: Scaling of the number of microstates with the number of ionizable residues for a perfect polyampholyte.** Even with the exclusion of forbidden microstates, the number of microstates grows "exponentially" with the number of ionizable residues. The total number of possible microstates is $2^n$, whereas the total number of microstates after elimination of the forbidden microstates is $2^{(n/2+1)}$-1.

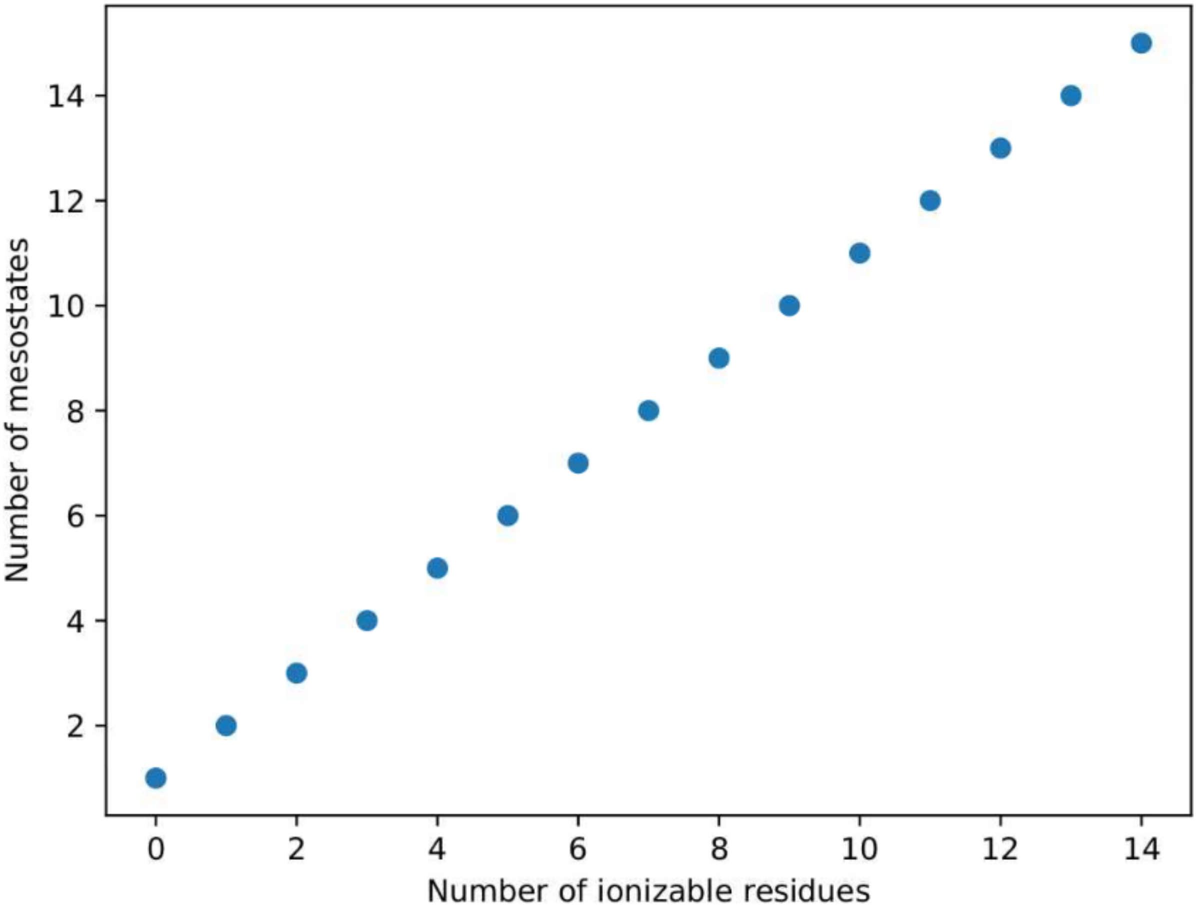


**Figure S2: The number of mesostates scales linearly with the number of ionizable residues.**

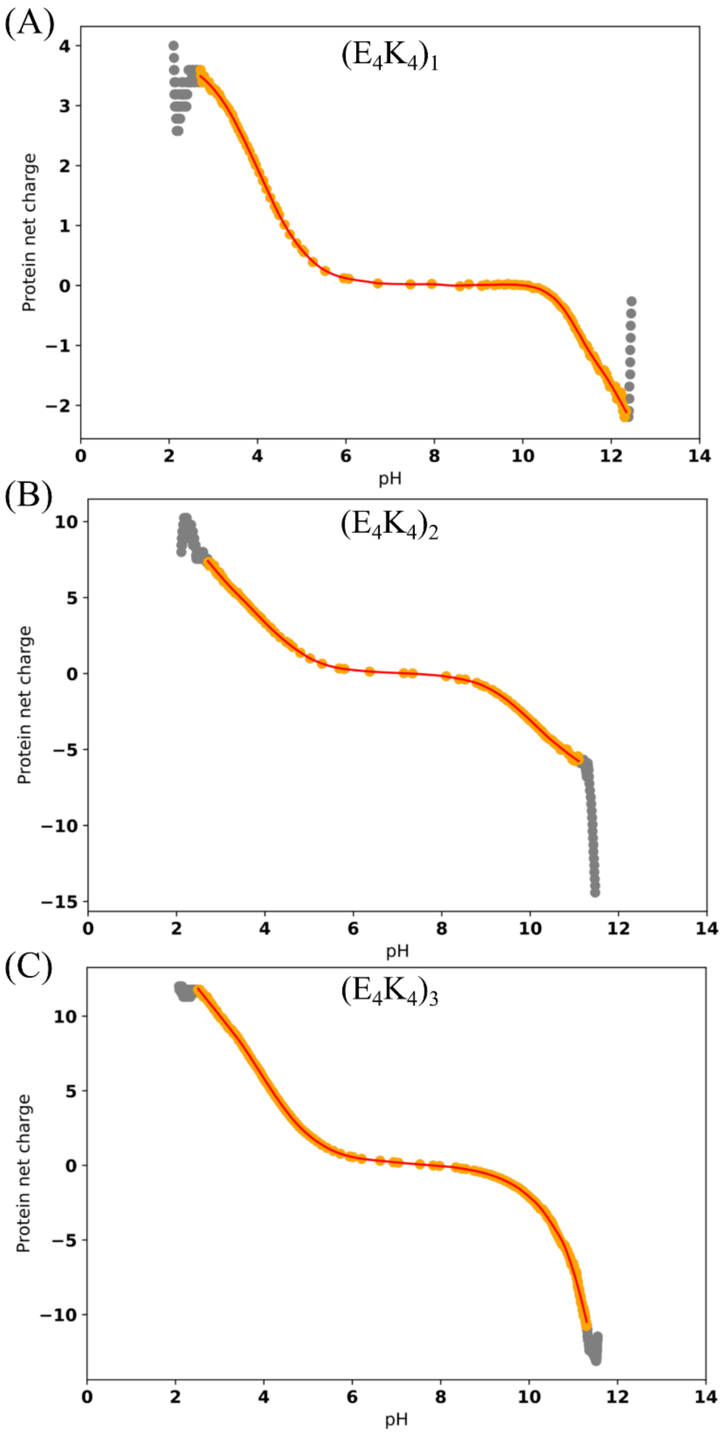


**Figure S3: Effect of smoothing on the data.** Raw data are shown in orange dots and resulting smoothed curve red. Results are shown for the $(E_4K_4)_1$, $(E_4K_4)_2$, $(E_4K_4)_3$.

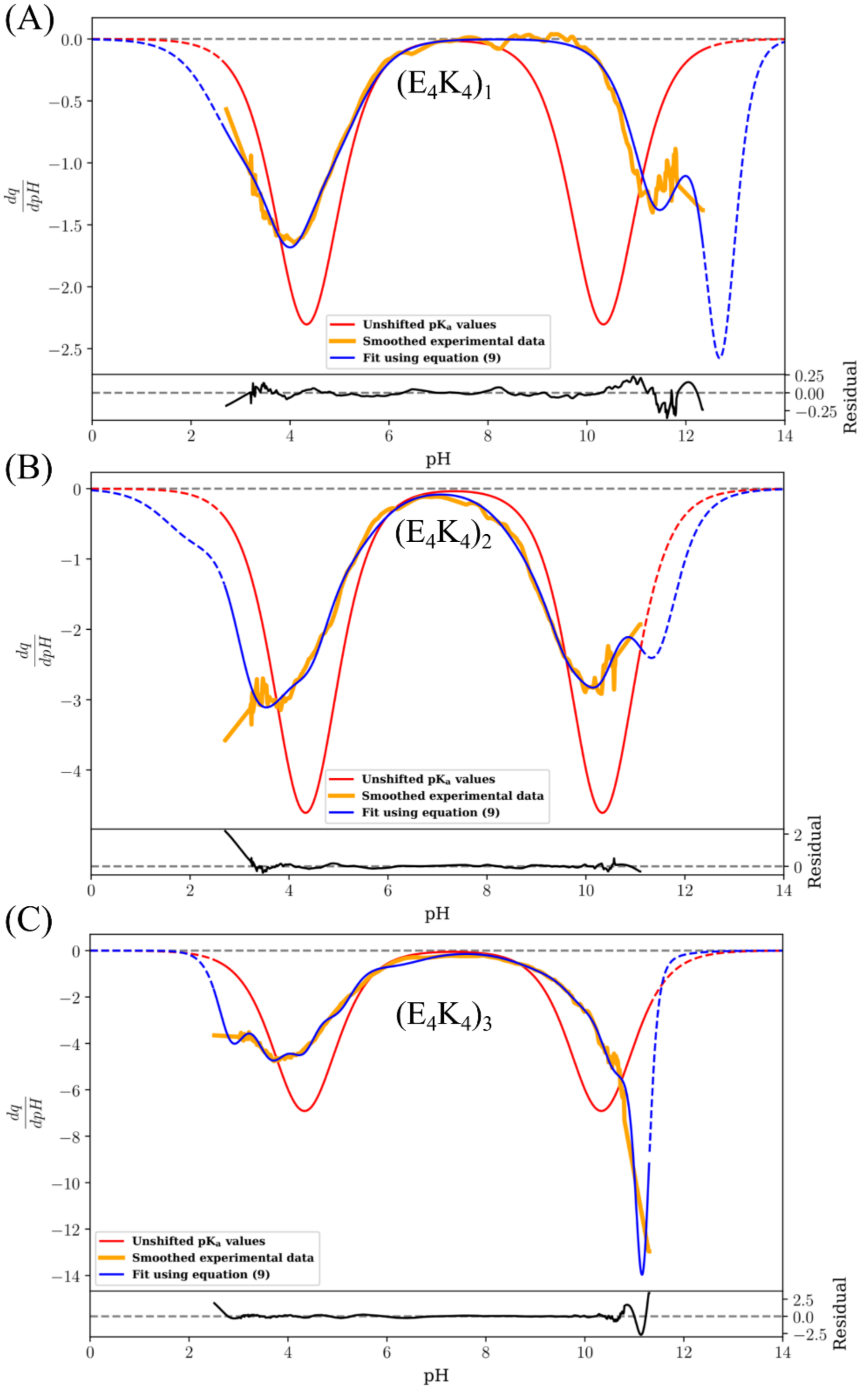


**Figure S4: Derivatives of each construct of the charge vs pH profiles.** The derivative of the smoothed data is shown in orange, the derivative of the fit is shown in blue. Each bottom panel quantifies the difference in between the two curves.

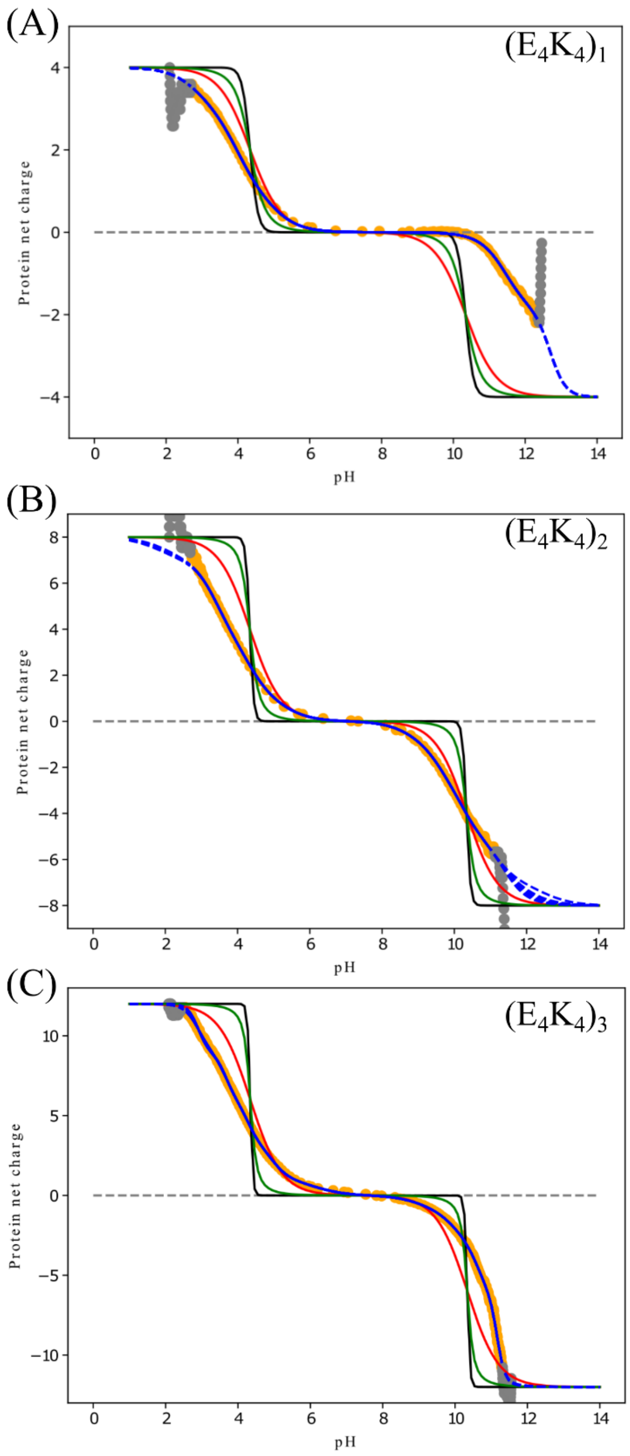


**Figure S5: Comparison of the 10 independent fits for each construct shown in blue.** The black curve shown the fixed charge model as in figure 4. The red curve shows the model calculated using model compound values and Equation (7). The green curve is the unshifted assumption ignoring sequence charge entropy.

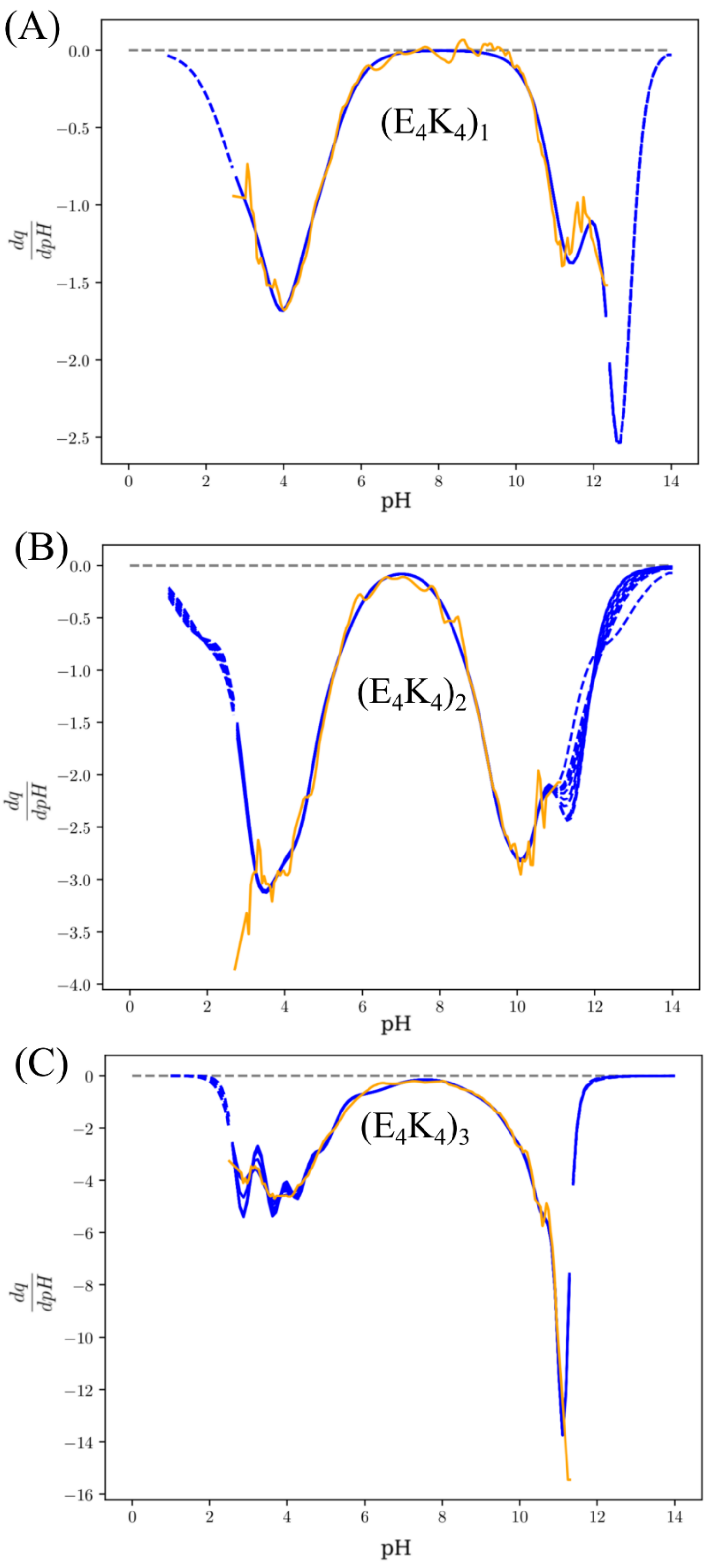


**Figure S6: Effect of the 10 independent fitting procedures for each construct on the derivative of the charge vs pH profile.** Color scheme is the same as in figure S2.

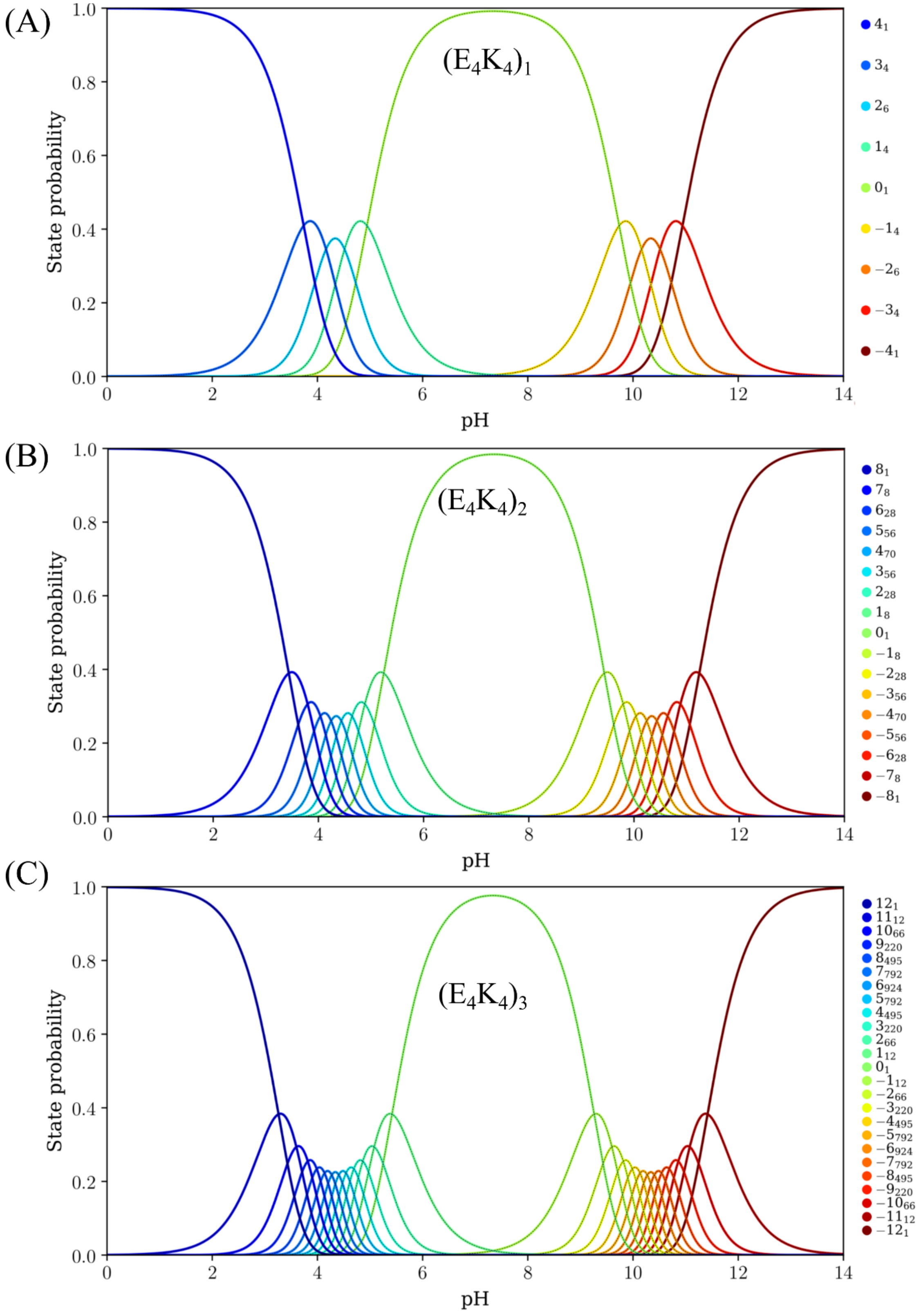


**Figure S7: Example of mesostates populations using the unshifted assumption and accounting for entropy as shown in Equation (7) for each of the constructs.**

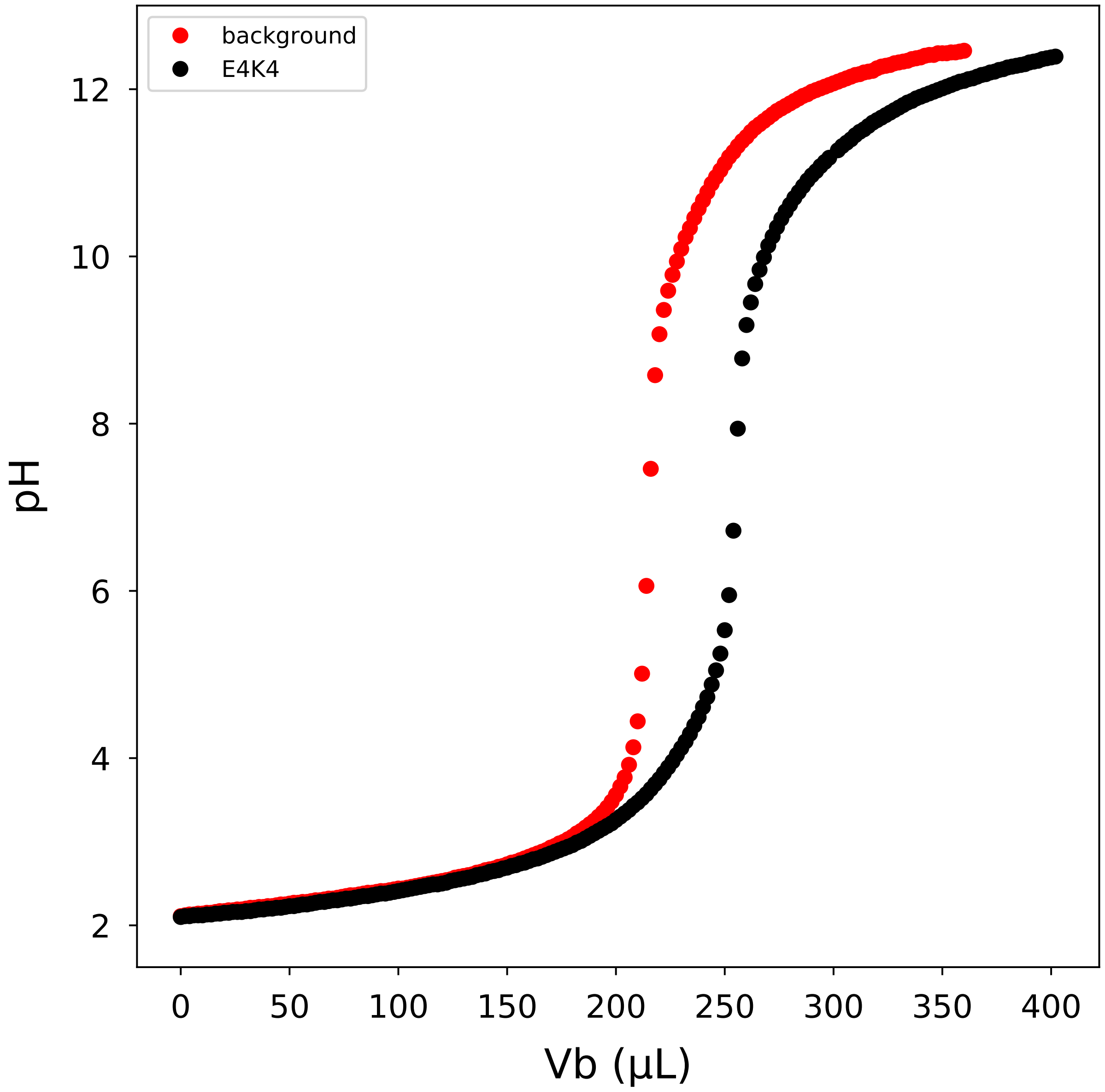


**Figure S8: Raw data from potentiometric measurments for $(E_4K_4)_1$.** The concentration of **$(E_4K_4)_1$** is 215 µM and the titrant (KOH) concentration is 48.3 mM. The background titration is shown in red, and the titration of the protein containing sample is shown in black.

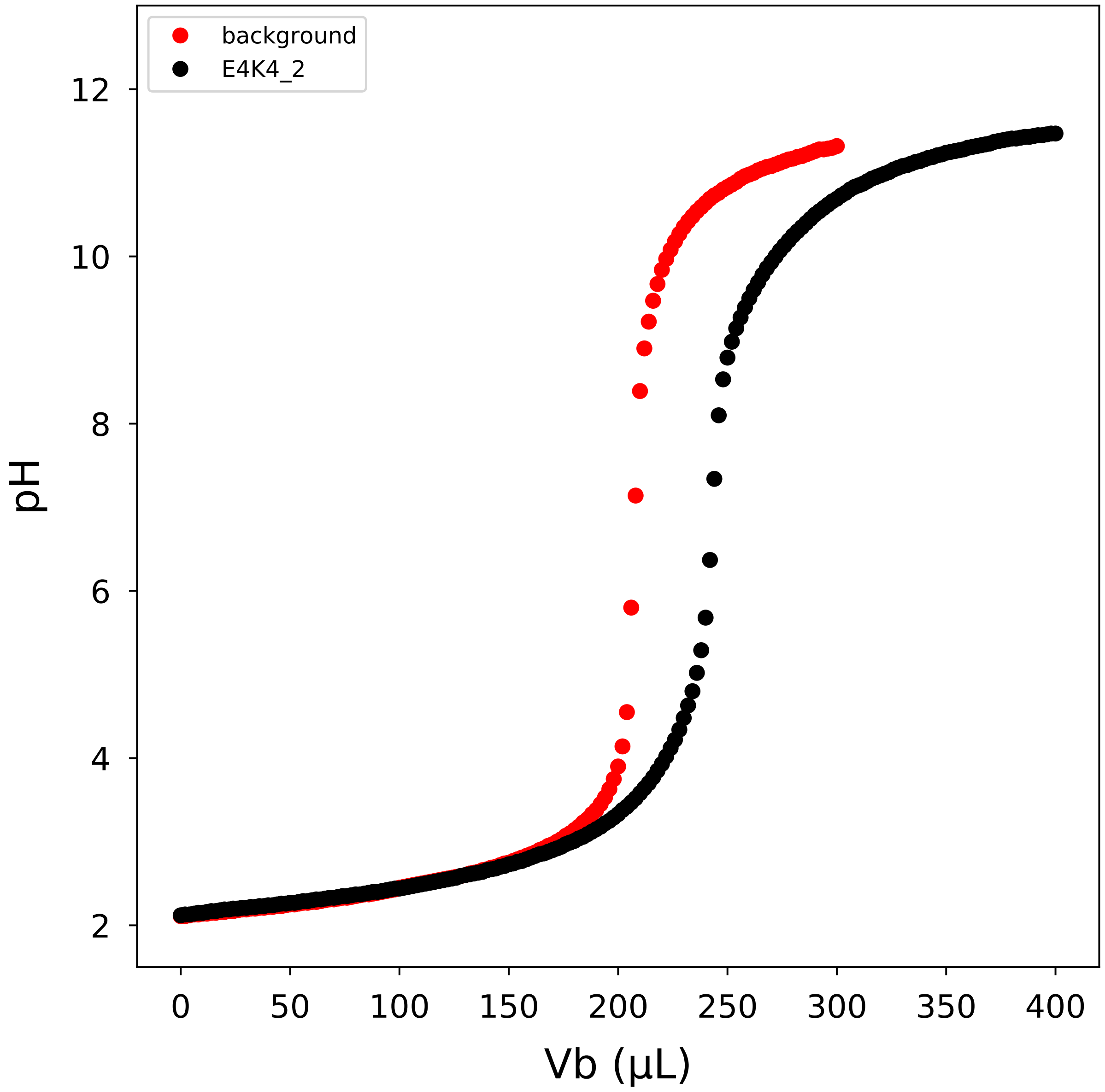


**Figure S9: Raw data from potentiometric measurments for $(E_4K_4)_2$.** The concentration of **$(E_4K_4)_2$** is 95 µM and the titrant (KOH) concentration is 56.0 mM. The background titration is shown in red, and the titration of the protein containing sample is shown in black.

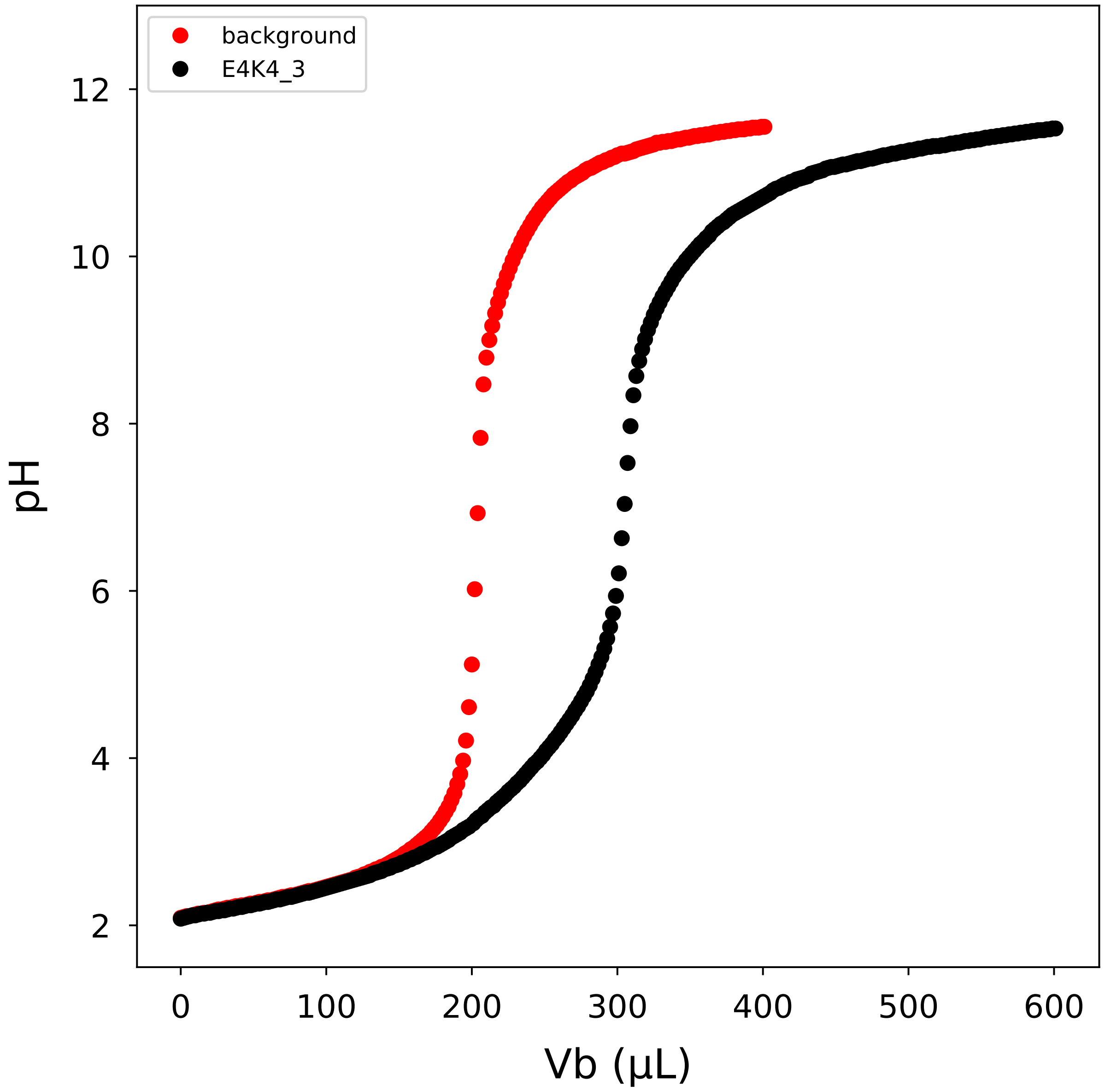


**Figure S10: Raw data from potentiometric measurments for $(E_4K_4)_3$.** The concentration of $(E_4K_4)_3$ is 180 µM and the titrant (KOH) concentration is 53.4 mM. The background titration is shown in red, and the titration of the protein containing sample is shown in black.

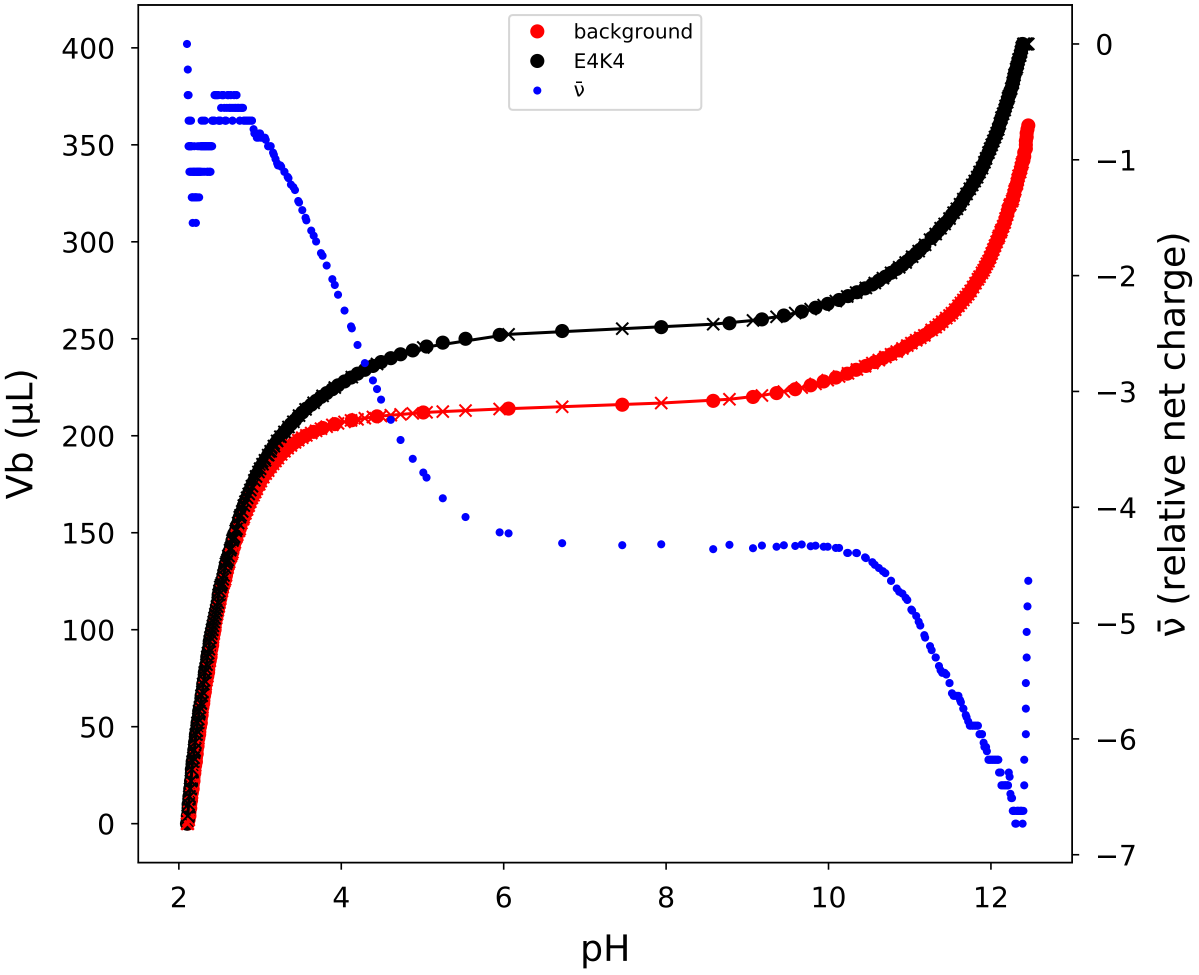


**Figure S11: Transposed titration data and relative net charge for $(E_4K_4)_1$.** Titration data were transposed, using the approach of Nozaki and Tanford, to facilitate comparison of the volume of KOH added in the background titration (red circles) vs. the peptide titration (black circles) at each pH value. Additional points (-x-) were calculated via linear interpolation between data points to facilitate subtraction of the two curves at each pH value. The moles of protons released per moles of peptide, which is equivalent to relative change in net charge (blue points), are calculated from the difference of the red and black data points (see main text for details).

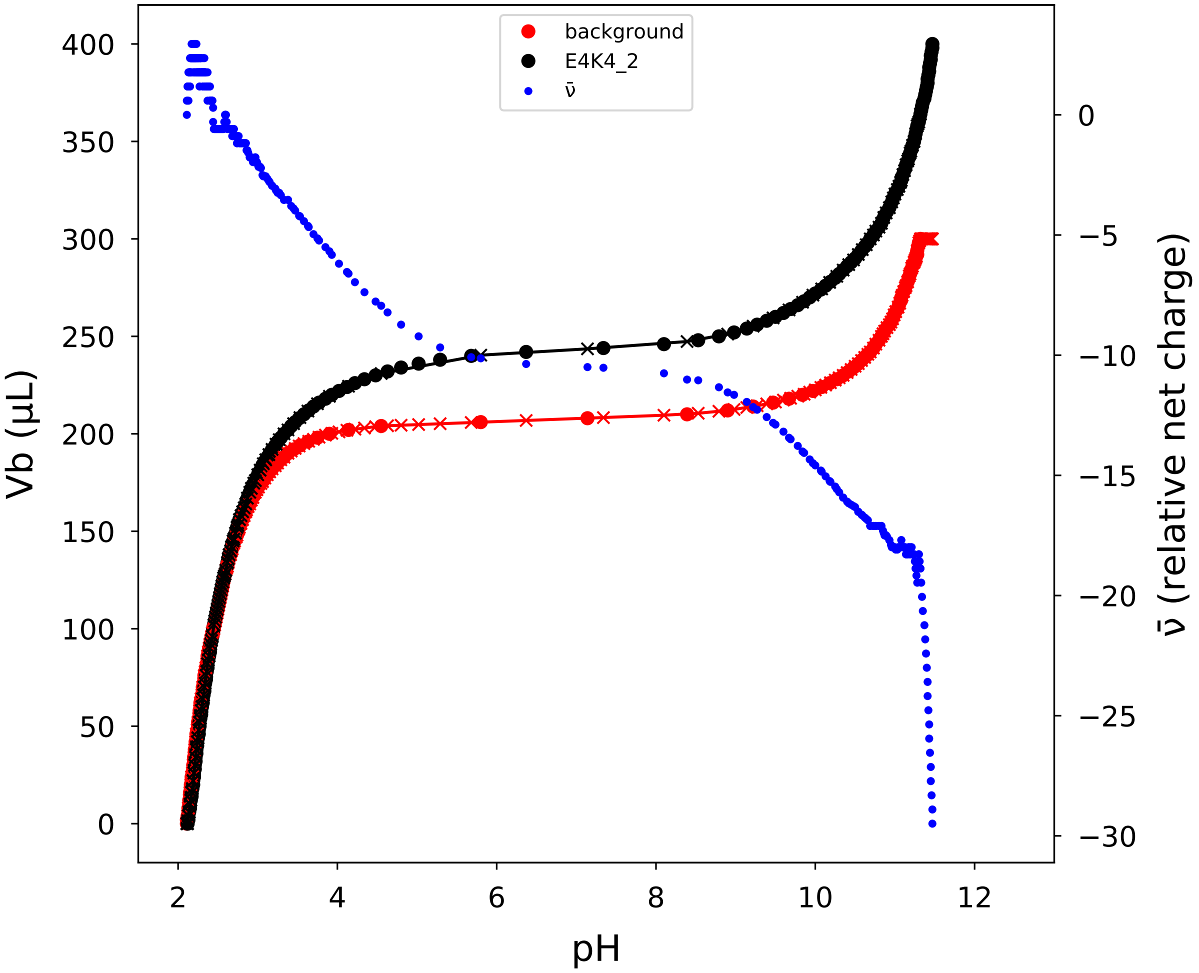


**Figure S12: Transposed titration data and relative net charge for $(E_4K_4)_2$.** Titration data were transposed, using the approach of Nozaki and Tanford, to facilitate comparison of the volume of KOH added in the background titration (red circles) vs. the peptide titration (black circles) at each pH value. Additional points (-x-) were calculated via linear interpolation between data points to facilitate subtraction of the two curves at each pH value. The moles of protons released per moles of peptide, which is equivalent to relative change in net charge (blue points), are calculated from the difference of the red and black data points (see main text for details).

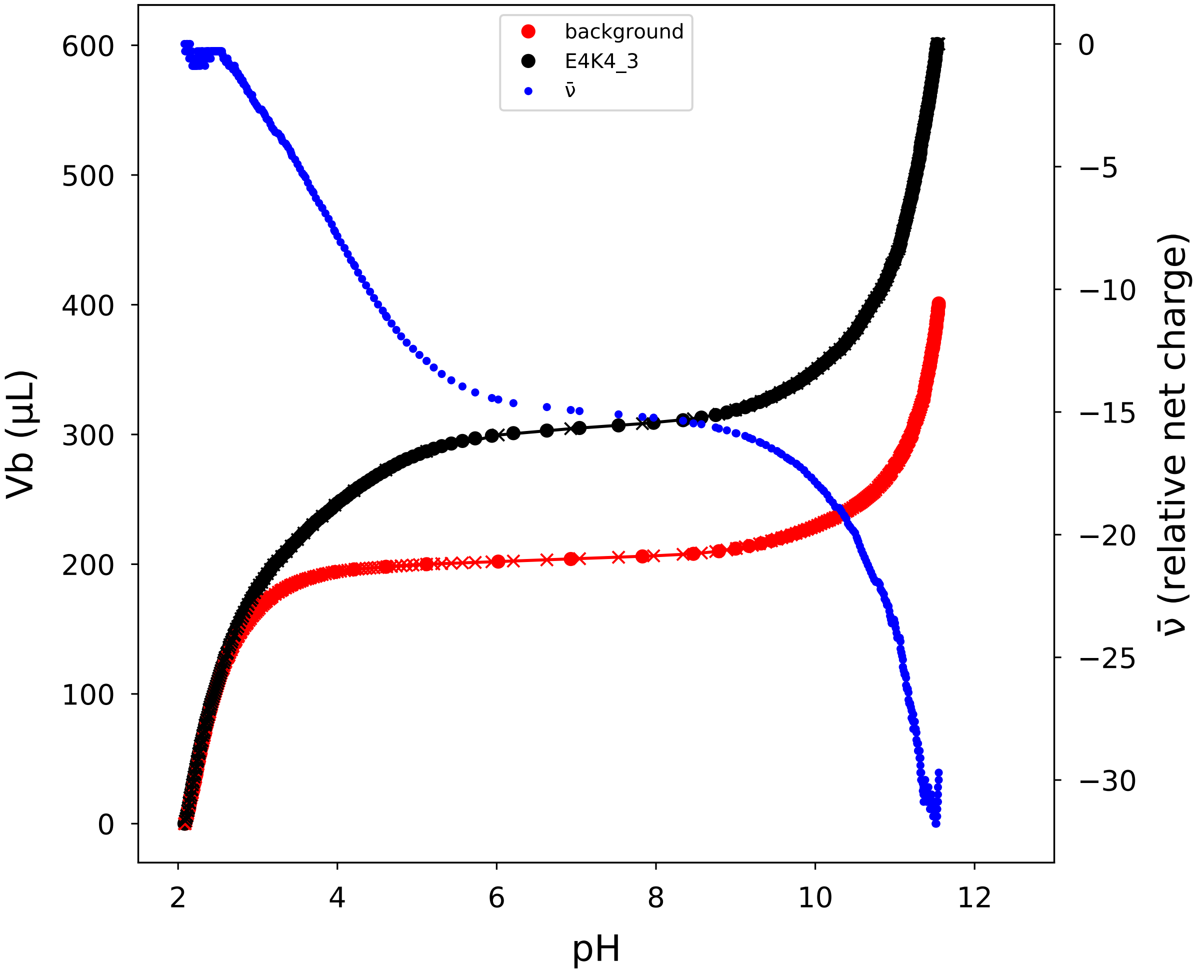


**Figure S13: Transposed titration data and relative net charge for $(E_4K_4)_3$.** Titration data were transposed, using the approach of Nozaki and Tanford, to facilitate comparison of the volume of KOH added in the background titration (red circles) vs. the peptide titration (black circles) at each pH value. Additional points (-x-) were calculated via linear interpolation between data points to facilitate subtraction of the two curves at each pH value. The moles of protons released per moles of peptide, which is equivalent to relative change in net charge (blue points), are calculated from the difference of the red and black data points (see main text for details).
